# Biochar Modulates Wheat Root Metabolome and Rhizosphere Microbiome in a Feedstock-dependent Manner

**DOI:** 10.1101/2024.07.17.604021

**Authors:** Hanyue Yang, Patricia Kerner, Xi Liang, Ethan Struhs, Amin Mirkouei, Yaqi You

## Abstract

**Background:** Biochar is a multifunctional soil conditioner capable of enhancing soil health and plant productivity, but the underlying mechanisms remain elusive. Here we tackled this question using wheat as a model plant and through the lens of the rhizosphere, a vital soil-plant interface continuum. We systematically examined the effects of four types of biochar (corn stover, cattle manure, pine sawdust, or wheat straw) applied at two rates (0.25% or 2.5%, w/w).

**Results:** Employing untargeted metabolomics and 16S rRNA gene sequencing, we revealed both common and unique modulating effects of the tested biochar treatments on wheat root metabolites and rhizosphere microbiome structure and functioning. Biochar modulated numerous metabolic pathways in wheat roots, where amino acid metabolism was the most common one, leading to cascade effects on the dynamics of a wide range of secondary metabolites, including many plant signaling molecules (e.g., flavonoid compounds, brassinosteroids) that are known to be involved in plant-microbe interactions. All biochar treatments increased rhizosphere microbial diversity, altered community composition, enhanced microbial interactions, and resulted in functional changes. Increased Burkholderiales (denitrifying bacteria) abundance and decreased Thermoplasmata (archaeal methanogens) abundance could explain biochar’s widely reported effects on nitrous oxide and methane mitigation, respectively. Biochar enhanced positive correlations among microbes and network complexity, particularly modularity, suggesting local adaptation through mutualism and/or synergism and the formation of modules of functionally interrelated taxa. A large number of diverse keystone taxa from both dominant and non-dominant phyla emerged after biochar treatments, including those known to be involved in methane, nitrogen, and sulfur cycling. Besides common alterations, treatment-specific alterations also occurred, and biochar type (i.e., feedstock choice) exerted greater influence than application rate. Wheat biochar applied at a 0.25% rate showed the strongest and distinct modulating effects, resulting in orchestrated changes in both root metabolites and rhizosphere microbiome, especially those relevant to plant-microbe interactions and likely beneficial to the host plant (e.g., upregulated biosynthesis of zeatin and down-regulated limonene degradation).

**Conclusions:** Our work contributes to a mechanistic understanding of how biochar modulates the soil-plant continuum and provides new insights into the potential of top-down rhizosphere microbiome engineering through biochar-based reprogramming of root-microbe interactions.

## Background

The global population is projected to reach ∼9.6 billion by 2050 and the food demand is forecasted to increase by 70-100%, with crop production estimated to rise by 80% from 2030 to 2050 (1-3). Soil produces a majority of human food and is essential for food security (4). Agricultural intensification has left unprecedented footprints on the earth causing soil quality degradation and biodiversity loss, water pollution due to nutrient leaching, increased greenhouse gas (GHG) emissions, among others (5). It is imperative to develop and implement sustainable practices to alleviate these deleterious outcomes while maintaining and improving crop yields. Biochar has long been proposed as a multifunctional soil enhancer (6). Numerous studies have shown that biochar amendment can simultaneously improve soil health, enhance crop productivity and resistance/resilience to stress, reduce nutrient leaching, and mitigate GHG emissions (6, 7). However, depending on the feedstock material and the production condition, biochar varies largely in physicochemical properties and thus amendment effects (8-10). Further, biological mechanisms underlying biochar’s beneficial effects are still unclear, which is critical for precision biochar production and application to elicit the desired benefits (8).

Soil is a living system where diverse microbes together mediate soil functions such as nutrient cycling and support plant development and health. Studies have shown that biochar amendment can modulate the bulk soil microbiome, changing microbial abundance, diversity, and community structure and function (11). A majority of previous studies attribute the phenomena to biochar-induced soil physicochemical improvements (e.g., water-holding capacity, acidity, nutrients, organic matter, toxic substances), which in turn enhance the abundance of plant-beneficial microbes while suppressing harmful ones (12). Few studies have explored biochar-mediated shifts in the rhizosphere, which may better explain biochar-induced benefits to the soil-plant continuum (8).

The rhizosphere is a thin layer of soil surrounding a plant’s root system and represents one of the most complex ecosystems on earth (13). This narrow zone encompasses a gradient of chemical and biological attributes that are shaped by entangled webs of interactions between the plant and its surrounding soil (14). Specifically, the rhizosphere microbiome is not a random assemblage but a result of long-term plant-microbiome co-evolution in the particular environment, and forms holobionts with the plant host (15, 16). Over the lifecycle of the plant host, the rhizosphere microbiome is dynamically recruited and assembled by the plant using root exudates and in turn provides benefits, including growth promotion, stress control, and defense against pathogens and pests, to the plant through direct and indirect mechanisms (14, 17). Yet it is unclear whether biochar can modulate the rhizosphere and thus benefit the plant host. Nor is it clear whether biochar characteristics or amendment practices can influence biochar effects on the rhizosphere.

This study aimed to use wheat as a model plant to gain a mechanistic understanding of how biochar amendment modulates the plant rhizosphere and how two management factors, feedstock choice and application rate, affect such rhizosphere modulation. Specifically, we asked: (1) Does biochar alter the root metabolome and the rhizosphere microbiome? (2) If so, how does feedstock choice or applicate rate influence rhizosphere alterations? (3) What root-microbe interactions are responsible for biochar-induced rhizosphere alterations? Addressing these questions contributes to a general understanding of the soil-plant continuum in response to a common sustainable agricultural practice. Further, rhizosphere microbiome engineering is an emerging tool that holds the promise of improving plant health by promoting beneficial plant-microbe interactions (18-21). Biochar amendment represents a top-down *in situ* manipulation approach and findings from this study can shed light on future targeted rhizosphere microbiome engineering.

## Methods

### Biochar production and plant growth experiment

Biochar was generated using four types of feedstocks (i.e., corn stover, cattle manure, pine sawdust, and wheat straw) by pyrolysis of the biomass feedstock at 350 ℃ under anaerobic conditions for 30 minutes like before (22). The resulting biochar was analyzed using a variety of methods to characterize physicochemical properties: Scanning electron microscopy (SEM) for microscopic structure (Fig S19), energy-dispersive X-ray spectroscopy (EDS) for surface elemental composition, and Boehm Titration for three oxygen-containing functional groups (acidic carboxyl, lactone, and phenolic group) on the biochar surface (23-26). In addition, biochar was analyzed at the Environmental Analytical Laboratory of Brigham Young University (Provo, Utah, USA) using conventional methods for soil to measure pH, major and trace elements, electrical conductivity, and cation exchange capacity (Additional file 8) (also see the Supplementary Information).

We chose bread wheat (*Triticum aestivum* L. cv. SY Ovation) as the model plant because it is one of the major sources of food for the world and the third largest crop grown in the United States. Wheat seeds were sterilized with 50% chlorox for 10 min, rinsed five times with sterile ultrapure water, and placed on moistened paper in sterile Petri dishes at 4 °C for germination (27). After germination, seedlings of similar size were sown in a pot (8.9 × 8.9 × 12.7 cm, L × W × H) filled with soil (coarse sandy loam, mixed, super active cation-exchange activity, mesic Xeric Haplocalcids) collected from an agricultural field near Aberdeen, Idaho, USA (soil property in Additional file 8). The soil was air-dried, crushed and sieved (2 mm mesh), homogenized, and stored at 4 °C. Before the pot experiments, the soil was placed at room temperature for two weeks to reactivate native microbial communities, completely mixed with biochar, and added to the pots to achieve the following treatments: (1) no biochar control, (2) 0.25% (w/w) corn biochar (C0.25), (3) 0.25% manure biochar (M0.25), (4) 0.25% pine biochar (P0.25), (5) 0.25% wheat biochar (W0.25), (6) 2.5% corn biochar (C2.5), (7) 2.5% manure biochar (M2.5), (8) 2.5% pine biochar (P2.5), and (9) 2.5% wheat biochar (W2.5). These two rates were chosen based on common application scenarios (8).

All treatments had four replicates, except for wheat biochar which had three replicates. The plants were cultivated in growth chambers (FXC-19, BioChambers) for 15 days: 50% humidity, 14 hours in light at 25 °C, 10 hours in dark at 15 °C, and watered once every two days with sterile ultrapure water.

### Collection of plant roots and ectorhizosphere soils

Sample processing was similar to our previous study (28). Each plant was carefully removed from the pot using aseptic techniques. The shoot system was separated from the root system using pre-sterilized scissors. Loosely bound soil was removed from the roots by shaking and using a sterilized spatula. The remaining roots with tightly bound soil (2–5 mm thick) were carefully transferred into a sterile 50 mL centrifuge tube, immediately frozen in liquid nitrogen (N), and stored at -80 °C until further processing as recommended earlier (29). To process rhizosphere samples, 15 mL autoclaved, cold 1× phosphate buffered saline (pH 7.4) and 15 μL Triton X-100 (0.22 μm filtered) were added to each 50 mL centrifuge tube and the tube vortexed gently without breaking roots apart. Surface-clean roots were aseptically transferred to a new sterile 50 mL centrifuge tube, immediately frozen in liquid N, and stored at -80 °C until metabolite analysis. The rhizosphere soil suspension was centrifuged at 4 °C, 6000 ×g for 10 min and the supernatant was pipetted out. This step was repeated three times to completely remove the supernatant. The soil pellet was stored at -80 °C until DNA extraction.

### Root metabolite extraction and metabolomic analysis

Endogenous metabolites were extracted as before with some modifications (30, 31). Briefly, surface-clean roots (n = 34) were pulverized in a 1.5 ml microcentrifuge tube using a Mini Bead Mill homogenizer (VWR International). A methanol-water mixture (80:20 v/v) was added to the tube at 1 mL per 100 mg wet root material. The tube was vortexed for 15 min, sonicated for 5 min, and centrifuged at 10,000 ×g for 5 min at room temperature. The resulting extracts were filtered through 0.45 µm micro-centrifugal PVDF filters and stored at -80 °C until untargeted metabolomics analysis.

Liquid chromatography with tandem mass spectrometry (LC-MS/MS) was performed on a Dionex UltiMate 3000 LC system (Thermo Scientific) coupled with a maXis electrospray ionization quadrupole time-of-flight (ESI Q-TOF) high-resolution MS system (Bruker Corporation) similarly to others (32). Samples of 10 µl were injected onto a Waters XTerra MS C18 column (5 µm, 2.1×150 mm) operated at 30 °C. The mobile phase consisted of 0.1% (v/v) formic acid in ultrapure water (solvent A) and 0.1% formic acid in acetonitrile (solvent B). The elution gradient began at 5% solvent B for 2 minutes and increased linearly to 80% solvent B over 43 minutes, after which the column was washed with 80% solvent B for 5 minutes and re-equilibrated for 10 minutes before the next injection. The flow rate was maintained at 200 µL/min. The MS conditions were as follows: ESI in positive ion mode with a spray voltage of 4000V, endplate offset of -500V, nebulizer gas pressure of 1.2 bar, dry gas flow rate of 8.0 L/min, dry gas temperature of 200 °C, and mass range of 20-800 m/z. Automated multiple reaction monitoring was performed, and auto MS/MS data acquisition used settings similar to others: precursor ions were acquired using a cycle time of 3.0 seconds for 3 ions; mass range was set to 20-800 m/z; ion polarity was positive; active exclusion was set to exclude after 3 spectra, release after 1.0 minute, and reconsider precursor if current intensity / previous intensity equaled 5× (32-34).

### Rhizosphere soil DNA extraction and 16S amplicon sequencing

Metagenomic DNA was extracted from ectorhizosphere soils (n = 34) using the PowerSoil Pro Kit (QIAGEN) following the manufacturer’s protocol. A PowerLyzer homogenizer (QIAGEN) was used to maximize extraction efficiency. Extracted DNA was further purified using the ethanol precipitation protocol (28). The resulting DNA was assessed on a NanoDrop 1000 spectrophotometer and by electrophoresis. 16S rRNA gene sequencing was performed as before(28). The V4 region of the prokaryotic 16S rRNA gene was amplified with forward barcoded 515F primer (5’-3’: GTGYCAGCMGCCGCGGTAA), 806R primer (5’-3’: GGACTACNVGGGTWTCTAAT), and Platinum Hot Start PCR Master Mix (Invitrogen) in a single PCR reaction (35-37). Libraries were cleaned up with magnetic beads (HighPrep PCR, MagBio Genomics), quantified on a CFX96 Touch Real-Time PCR System (Bio-Rad Laboratories) with a NEBNext Library Quant Kit for Illumina (New England BioLabs), assessed by fragment analysis on a 5200 Fragment Analyzer (Agilent Technologies), quantified on a Qubit fluorometer (Invitrogen), and pooled in equimolar concentrations for sequencing on an Illumina MiSeq platform (2×250 bp).

## Bioinformatics

### Metabolic data processing and analysis

Instrument-specific spectrum data were converted to the generic mzXML format using the msConvert toolkit of ProteoWizard (38, 39) and then processed using four platforms (MetaboAnalyst 5.0, XCMS, OpenMS, MZmine3) under default parameter settings according to each platform’s guidelines (40-44). The three online platforms (MetaboAnalyst 5.0, XCMS, OpenMS) have similar steps in general, including peak picking, peak alignment, gap filling, and peak annotation, but use different databases and algorithms. In MetaboAnalyst 5.0, we used the algorithm centWave (continuous wavelet transformation) and LOESS (local nonlinear regression without internal standards) for peak picking and alignment, respectively (40). In XCMS, we used the algorithm centWave and obiwarp (all peaks aligned toward a center sample) for peak picking and alignment, respectively (44). In OpenMS, whose workflow differs from the two online platforms (Fig S20), we used default parameter settings (42). In MZmine3, we used the ADAP chromatogram builder module with smoothing for feature detection, the Join aligner module (all peaks aligned through a match score based on specified m/z and retention time tolerances) for feature alignment, and the Peak finder module (“back-filling” approach using specified m/z and retention time tolerances) for gap-filling (43).

Feature tables from the four platforms were separately input to MetaboAnalyst 5.0 for functional analysis with the mummichog algorithm (45): positive ion mode (mass tolerance = 5ppm); data filtering with standard deviation (SD) set to 25% for feature tables with <1000 features (MetaboAnalyst 5.0, OpenMS, MZmine3) and 40% for the feature table with >1000 features (XCMS) according to the protocol (46); data pretreatment by median normalization and log transformation; mummichog v2.0 with a *p*-value cutoff of 0.05 for pathway activity prediction based on the Arabidopsis thaliana (*thale cress*) library; enriched pathway exploration within the KEGG global metabolic network; for missing values in two-group comparisons, compounds exclusive to one group and present in <50% of the samples were removed. Compound tables and enriched pathway tables generated from individual feature tables were exported and merged for (without redundancy) before downstream analysis (more details in the Supporting Information). Compound names were converted to the International Union of Pure and Applied Chemistry (IUPAC) names and classified to the “Class” level of the purely structure-based chemical taxonomy (ChemOnt) using ClassyFire (47). For fold change (FC) calculation, compound intensity values were transformed by adding 0.001 to the original values.

Principal coordinates analysis (PCoA) was performed using the R package “vegan” (Jaccard similarity) to explore total variance in root metabolites after varying treatments (48). To identify metabolites most responsible for specific treatments, we employed canonical discriminant analysis (CDA, also called linear discriminant analysis) to ∼600 compounds that were significantly influenced by any treatment as compared to control (FC > 1.5 and *p* < 0.05) (Additional file 1). Unlike principal components analysis (PCA), CDA generates latent discriminant functions to maximize between-class objective dispersion and when the result is visualized as a scatter plot, discriminant functions serve as axes (49). Since the first canonical axis (CD1) of the resulting CDA explained a vast majority (42.16%) of the total variance among biochar-influenced metabolites (Additional file 9), we extracted the absolute CD1 values of these metabolites and fitted them to a two-parameter log-normal distribution. Metabolites that had CD1 values above the 90th percentile of the fitted distribution were considered common biochar-influenced metabolites (50, 51).

### 16S amplicon sequencing data processing and analysis

Paired-end reads were processed using QIIME2 as before (28, 52). Briefly, raw reads were trimmed at both ends when the Phred score dropped below 33 and denoised with DADA2 to identify amplicon sequence variants (ASVs) (53). Rare ASVs (<5 occurrences among all the samples or only present in one sample) were filtered out to avoid sequencing errors and artifacts. Taxonomy was assigned using a naive Bayes machine-learning classifier trained for the 515F-806R V4 region against the SILVA database (release 138) (54, 55). Unassigned reads and reads associated with Eukarya, mitochondria, and chloroplast were removed prior to downstream analysis with the R package “phyloseq” (56).

Alpha diversity (Shannon, Simpson, ACE, and Chao1) was assessed at the ASV level after rarefaction. Beta diversity was estimated using Bray-Curtis of log transformed ASV abundance and weighted UniFrac distances of ASV abundance, and visualized using PCoA (57). We applied linear discriminant analysis (LDA) of effect size (LEfSe) on ASV relative abundance to identify differential taxa most likely to explain differences between samples, where the threshold on the logarithmic LDA absolute value was set to 2 and the alpha values for the ANOVA and Wilcoxon tests were set to 0.05 (58).

PICRUSt2 was used to infer metagenomic functional contents (59). The abundance of gene families was inferred based on KEGG orthologs (KOs) and Enzyme Commission (EC) numbers. The abundance of the pathways was inferred based on the MetaCyc database through structured mappings of EC gene families. The nearest-sequenced taxon index (NSTI) was calculated and any ASV with NSTI > 2 was excluded from the output. PICRUSt2 predictions were subjected to differential abundance analysis using the R package ggpicrust2 (60).

### Network analysis of multi-omics data

For the root metabolite network, we calculated pairwise Pearson correlation (Benjamini-Hochberg (BH) false discovery rate (FDR) procedure) using the R package “WGCNA” (61). Significant correlations (*r* > 0.9 and adjusted *p* < 0.01) were retained. Network topology analysis was conducted using the “igraph” (62). Node degrees were fitted to power-law distribution, and nodes with degrees above the 95% quantile were considered highly connected hub nodes (63).

For the microbial network, the pairwise correlation was calculated using SparCC, an approach more appropriate for compositional data, with *p*-value determined by bootstrapping (1000 times) (64, 65). Significant correlations (*p* < 0.05) were retained and analyzed for network topology. We identified hub nodes as nodes above the 95% quantile of the fitted log-normal distribution of three network centrality measures: degree, betweenness, and closeness (66).

Among all treatments in this study, W0.25 showed the strongest modulating effects on the root metabolome and the rhizosphere microbiome. Therefore, we explored correlations between significantly altered root metabolites and rhizosphere microbes whose predicted functions were relevant to root metabolites under control vs. W0.25. Microbe-metabolite co-occurrence networks were constructed using the data of microbial genera and root metabolites on the multi-omics analysis platform OmicsNet 2.0 (67). Metabolic potential of microbial genera and relevant proteins were predicted using the AGORA database of high-quality genome-scale metabolic models (potential score 0.8; excluding currency metabolites, universal metabolites, and metabolites without pathway annotation) (68). Microbial protein-root metabolite network was constructed based on all KEGG reactions (67). Because root metabolites rather than microbial metabolites were used here, the OmicsNet 2.0-predicted microbial pathways (FDR-adjusted *p* < 0.01) were mapped to the PICRUSt2-predicted microbial functions (FDR-adjusted *p* < 0.01). Eighteen pathways were retained, and the contributing microbial genera (nodes in the multi-omics networks) were retrieved (Additional file 7).

### General statistical analysis and data visualization

Statistical analysis was conducted in R (v 4.2.1) and Origin (v 2019b). Significant differences between treatments were tested using one-way analysis of variance (ANOVA) for biochar properties and wheat height, and the nonparametric Wilcoxon rank test or Kruskal-Wallis test for other comparisons such as microbial alpha diversity. Significant differences in microbial beta diversity were determined using permutational multivariate analysis of variance (PERMANOVA) (69). Cytoscape (v 3.9.1) was used for all network visualization. The R packages “pheatmap”, “VennDiagram”, and “ggplot2” were used for data visualization.

## Results

### Biochar modulated root metabolism of wheat plants

After fifteen days of pot experiments, 0.25% biochar treatments generally enhanced wheat height, except for manure biochar, but this effect was statistically insignificant. Amendment with 2.5% biochar, regardless of the biochar type, generally decreased wheat height and this effect was significant for manure biochar (Fig S1). Biochar significantly influenced the root metabolome of wheat (PERMANOVA: *p* = 0.001), affecting about 800 out of the 953 identified metabolites (Wilcoxon rank: *p* < 0.05) (Fig 1A and S2) (Additional file 1). Across biochar treatments, more metabolites were down-regulated than up-regulated, with feedstock (PERMANOVA: *p* = 0.001) and application rate (PERMANOVA: *p* = 0.042 for root glycans) being two influential factors. A majority of the observed metabolic alterations occurred to all biochar types: A total of 665 and 630 metabolites were responsive to 0.25% and 2.5% application rate, respectively (Fig 1B-C), and 597 metabolites were commonly influenced by all biochar types and application rates. A few of metabolites were only responsive to specific treatments: 8, 32, 24, and 58 metabolites were uniquely responsive to C0.25, M0.25, P0.25, and W0.25, respectively; 20, 31, 36, and 25 metabolites were uniquely responsive to C2.5, M2.5, P2.5, and W2.5, respectively.

**Fig 1.**
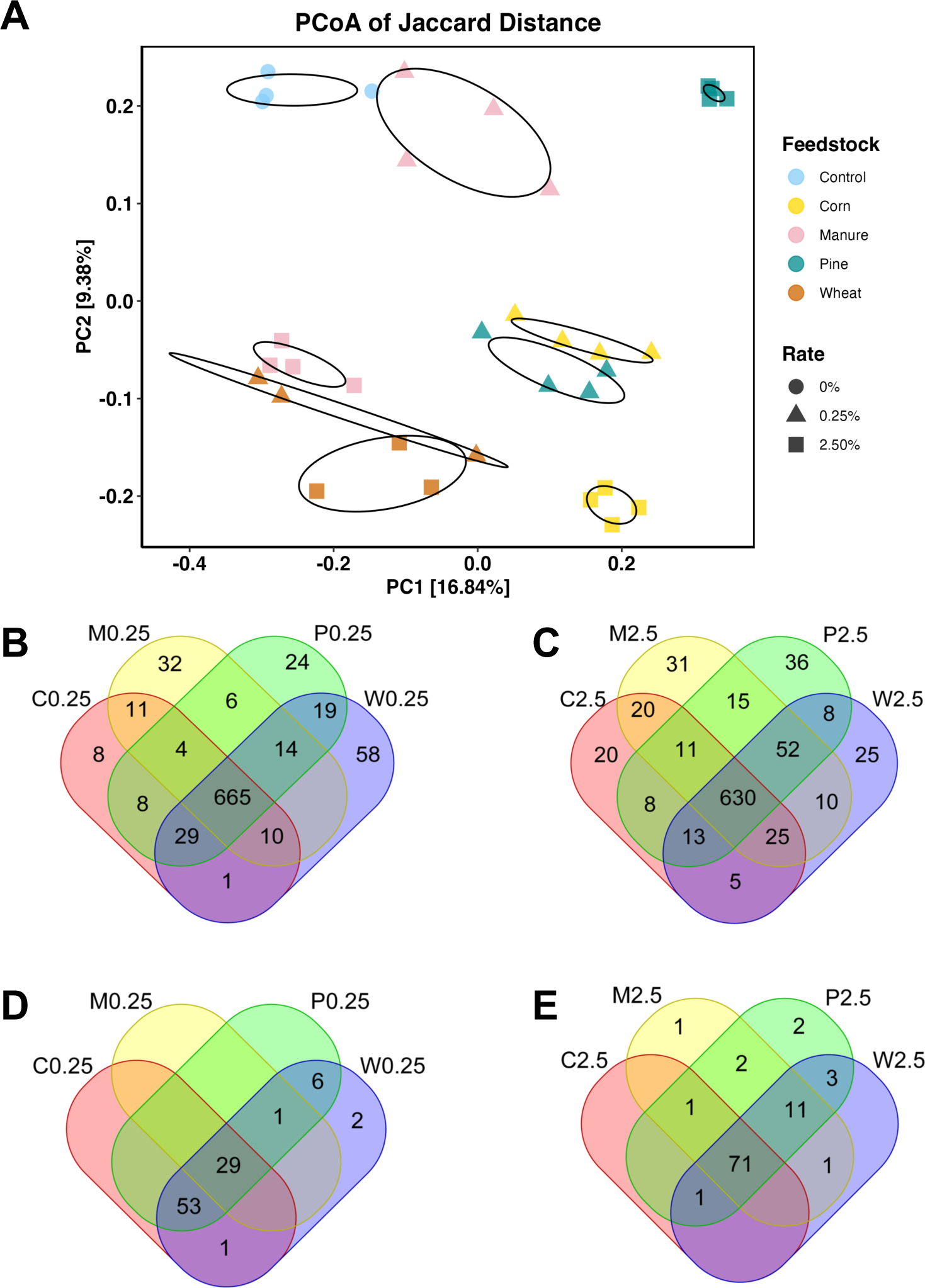
Biochar modulated the wheat root metabolome in a feedstock- and rate-dependent manner. (A) PCoA of detected root metabolites based on Jaccard distance. Ellipses represent 95% confidence. (B-E) Venn diagram showing the number of metabolites (B, C) and metabolic pathways (D, E) significantly altered. Biochar treatments are indicated by feedstock (C, corn stover; M, cattle manure; P, pine sawdust; W, wheat straw) and application rate (0.25% or 2.5%).

Across all treatments, 25 metabolic pathways from 8 KGEE pathway classes were commonly affected: carbohydrate metabolism (pentose and glucuronate interconversions), lipid metabolism (alpha-linolenic acid metabolism; steroid biosynthesis), nucleotide metabolism (pyrimidine metabolism), amino acid metabolism (alanine, aspartate and glutamate metabolism; cysteine and methionine metabolism; arginine and proline metabolism; phenylalanine metabolism; tryptophan metabolism; phenylalanine, tyrosine and tryptophan biosynthesis), glycan biosynthesis and metabolism (N-glycan biosynthesis); metabolism of cofactors and vitamins (folate biosynthesis; riboflavin metabolism; vitamin B6 metabolism; ubiquinone and other terpenoid-quinone biosynthesis); metabolism of terpenoids and polyketides (brassinosteroid biosynthesis; carotenoid biosynthesis; diterpenoid biosynthesis; sesquiterpenoid and triterpenoid biosynthesis); biosynthesis of other secondary metabolites (phenylpropanoid biosynthesis; flavonoid biosynthesis; flavone and flavonol biosynthesis; anthocyanin biosynthesis; glucosinolate biosynthesis), translation (aminoacyl-tRNA biosynthesis) (Fig 1D-E) (Additional file 2). Twenty-nine and 71 pathways were responsive to 0.25% and 2.5% biochar, respectively, regardless of the biochar type. Glycosaminoglycan degradation (glycan biosynthesis and metabolism class) was only responsive to W0.25; linoleic acid metabolism (lipid metabolism) and glycosphingolipid biosynthesis - ganglio series (glycan biosynthesis and metabolism) were unique to P2.5. Overall, W0.25 treatment manifested the strongest modulation of root metabolism, inducing unique alterations not seen in other treatments.

CDA pinpointed 44 discriminative metabolites across all treatments (Fig 2). Half of them were associated with 14 pathways, accounting for 56% of the 25 commonly influenced pathways. These metabolites fell into 13 ChemOnt classes, including 9 prenol lipids, 7 organooxygen compounds, 4 carboxylic acids and derivatives, 4 flavonoids, 3 steroids and steroid derivatives, 3 keto acids and derivatives, 2 fatty acyls, 2 indoles and derivatives, 1 imidazopyrimidines, 1 organonitrogen compound, 1 pteridines and derivatives, 1 pyrimidine nucleotides, among others. Significantly decreased metabolites across all treatments (mean FC ≤ 0.25) included 2 carboxylic acids and derivatives (cysteinylglycine, hydroxythreonine), 3 flavonoids (cyanidin 3-glucoside, leucodelphinidin, quercetin-3-O-rhamnoside-7-O-glucoside), 2 indoles and derivatives (indole-3-acetaldehyde oxime, indole-3-acetamide), 3 organooxygen compounds (5-aminopentanal, galabiose, glucotropaeolin), 1 pteridines and derivatives (5,10-methylene-THF), 1 purine nucleotides (deoxyadenosine), 2 steroids and steroid derivatives ((GlcNAc)2(Man)7(PP-Dol)1, 16-feruloyloxypalmitic_acid, glycineamideribotide), and other metabolites. Significantly increased metabolites (FC ≥ 1.5) only occurred to some treatments, including 1 carboxylic acids and derivatives (ornithine), 1 fatty acyls (2-isopropylmalic acid), 1 flavonoids (pinobanksin 5-[galactosyl-(1->4)-glucoside]), 1 keto acids and derivatives (2-dehydropantoate), 4 prenol lipids (abscisic_acid_glucose_ester, gibberellin_A15_open_lactone, cis-neoxanthin, gamma-tocopherol), 1 pyrimidine nucleotides (deoxycytidine).

**Fig 2.**
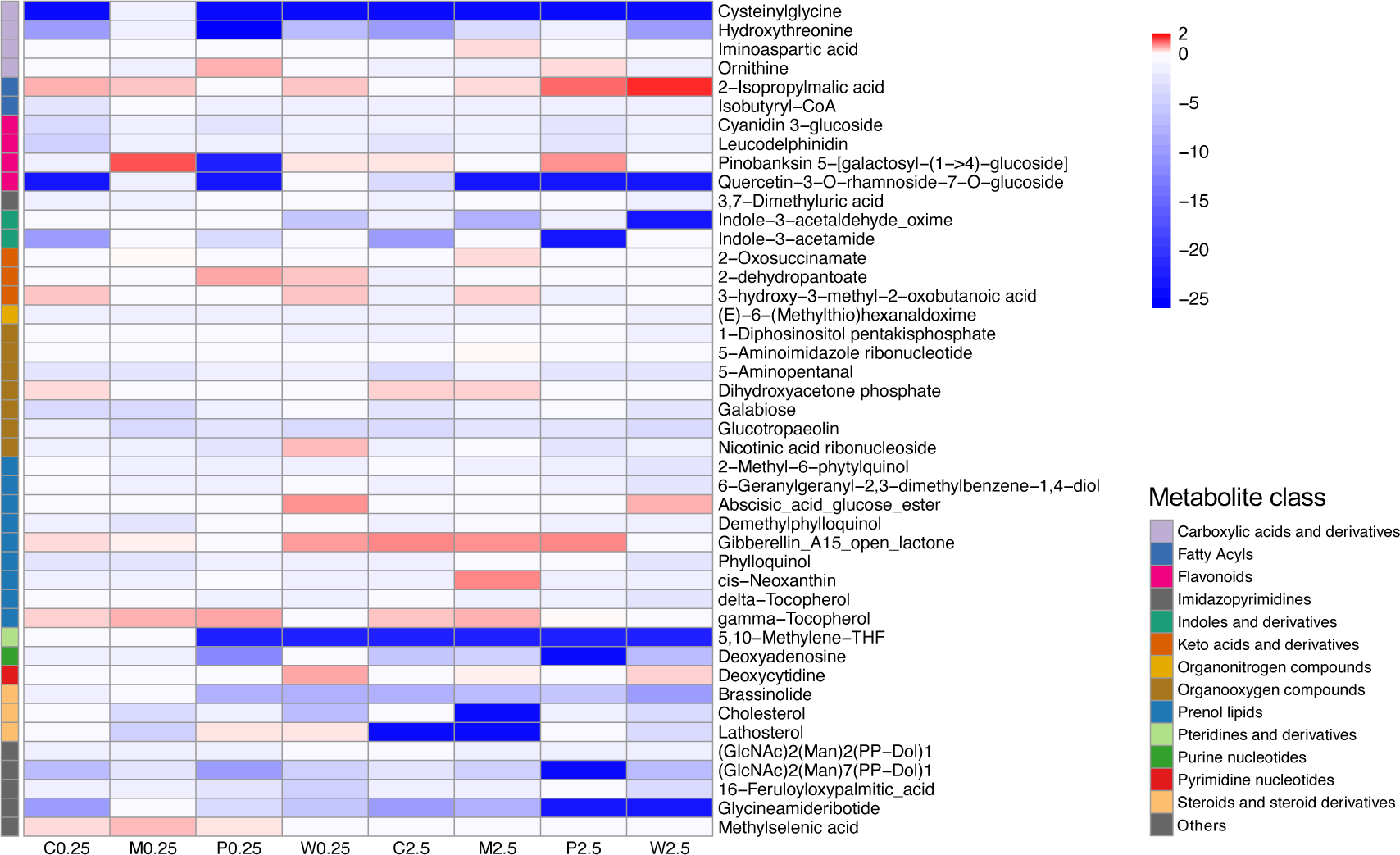
Heat map of forty-four most discriminative metabolites in biochar-treated wheat roots. Blue and red colors indicate normalized fold change compared to non-biochar controls. Individual metabolites are also grouped into major classes reflecting the relevant pathways.

Biochar regulation on root metabolism was also manifested by a general increase in average degree and centrality of the metabolite network (Table S1). Hub networks further revealed differential keystone metabolites in biochar-treated roots compared to the control, as well as across treatments (Fig 3) (Additional file 3). Notably, the metabolites responsive to specific treatments, as shown in Fig 1B-C were not keystone metabolites. The most drastic changes occurred to W0.25, followed by W2.5, where a large number of keystone metabolites (e.g., carboxylic acids and derivatives, flavonoids, indoles and derivatives, organic phosphoric acids and derivatives, phenols, steroids and steroid derivatives) not seen under other treatments emerged and connections of many metabolites substantially increased. While the C2.5 hub network also had new keystone metabolites compared to the control, its global connectivity was less prominent than the W0.25 or W2.5 hub network, and the whole network fell into several separate clusters.

**Fig 3.**
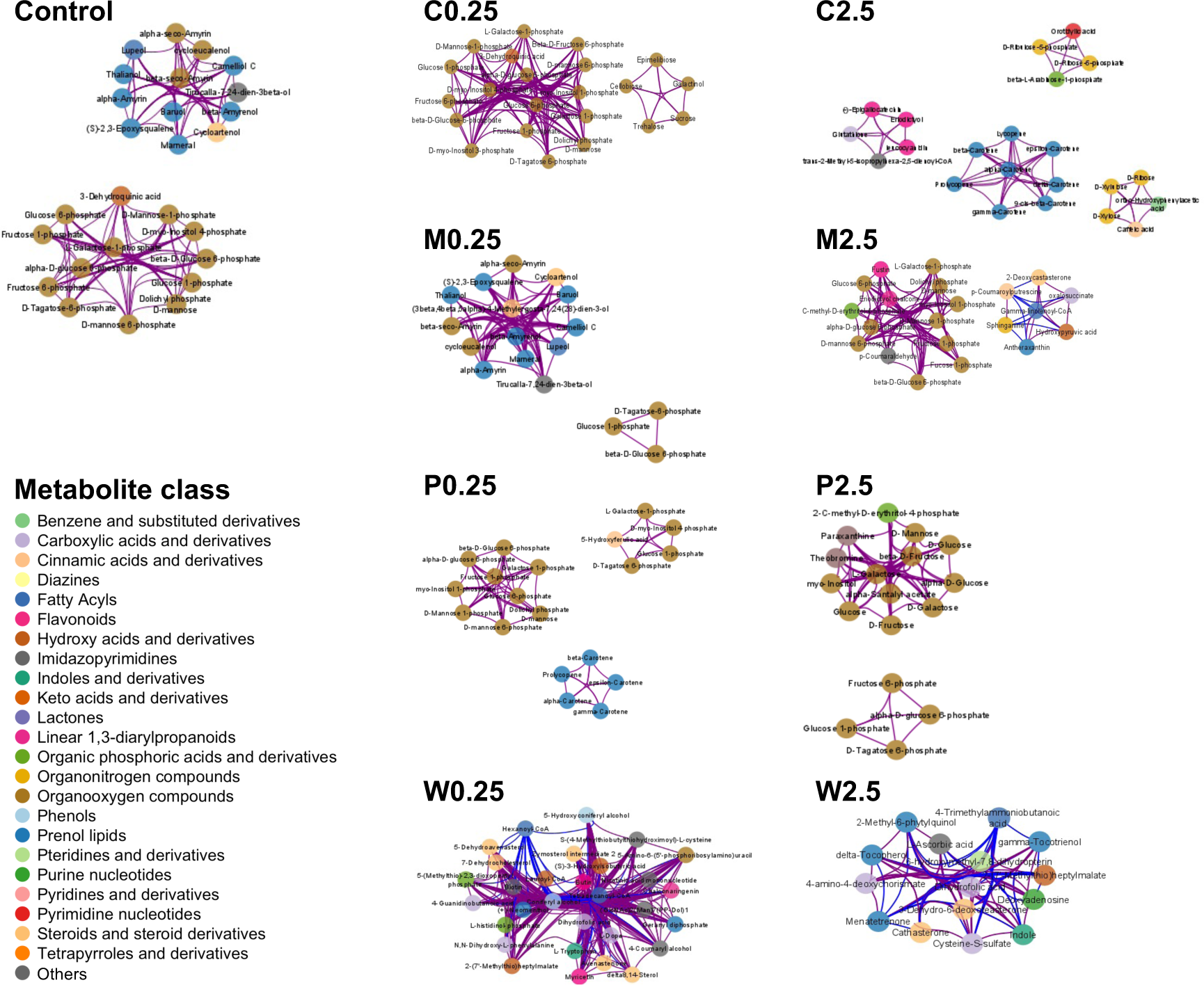
Hub network of keystone root metabolites in various treatments. Keystone metabolites were defined as nodes with the highest degrees (95% quantile of the fitted power-law distribution) in individual metabolite co-occurrence networks. Node colors indicate metabolite classes and metabolite names are labeled. Purple and blue edges indicate positive and negative correlations, respectively.

### Biochar reshaped the rhizosphere microbiome of wheat plants

In general, biochar increased microbial (archaea and bacteria) alpha diversity (Shannon, Simpson, ACE, Chao1). This effect was significant for Shannon diversity (Kruskal-Wallis: *p* = 0.028) (Fig S3). Regardless of the feedstock, 0.25% biochar increased Shannon diversity from 6.04 ± 0.46 (mean ± SD) to 6.30 – 6.63 (Kruskal-Wallis: *p* = 0.043) (Fig S4-S8). Biochar at the 2.5% application rate increased Shannon diversity to 6.15 – 6.43, less than 0.25% biochar, and the effect was insignificant (Kruskal-Wallis: *p* = 0.258) (Fig S4-S8).

Further, biochar treatments reshaped the rhizosphere microbiome as shown in the PCoA plot of UniFrac distance of ASVs (PERMANOVA: *p* < 0.001 for both feedstock and application rate) (Fig 4A). The influence of biochar feedstock was largely captured by the first coordinate, which explains 27.13% of the total variance. The influence of biochar application rate was captured by the second coordinate, which explains 8.85% of the total variance. Similar results were observed in the PCoA plot of Bray-Curtis distance of ASVs (PERMANOVA: *p* < 0.003 for both feedstock and application rate) (Fig S9).

**Fig 4.**
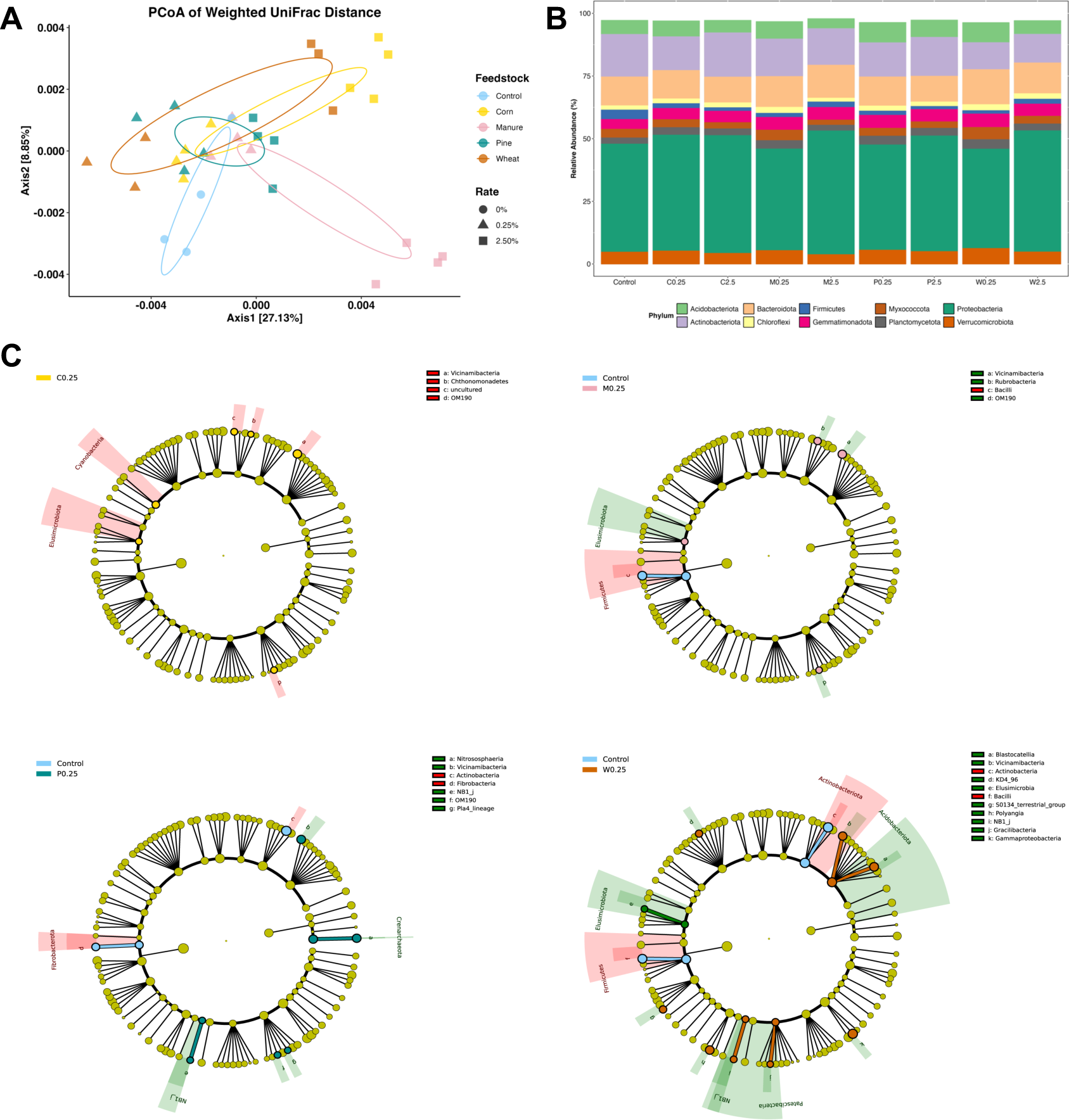
Biochar reshaped the rhizosphere microbiome of wheat plants. (A) PCoA of weighted UniFrac distance of ASVs. (B) Relative abundance of the top 10 microbial phyla. (C) Microbial taxa significantly influenced by four types of biochar applied at 0.25%. Only phyla and classes are shown in the LEfSe cladograms.

In the native wheat rhizosphere, Proteobacteria, Actinobacteriota, Bacteroidota, Acidobacteriota, Verrucomicrobiota, Gemmatimonadota, Firmicutes, Myxococcota, Planctomycetota, and Chloroflexi were the top 10 phyla (in descending order), comprising more than 95% of the microbiome (Fig 4B) and reflecting a core microbiome in the plant rhizosphere (14). Biochar influenced many microbial taxa, with feedstock and application rate being influential factors (Fig 4C and S10). At the phylum level, C0.25 enriched Cyanobacteria and Elusimicrobiota; M0.25 enriched Elusimicrobiota while suppressing Firmicutes; P0.25 enriched Proteobacteria (NB1-j under the Deltaproteobacteria class) and the archaeal phylum Crenarchaeota while suppressing Fibrobacterota; W0.25 enriched Acidobacteriota, Elusimicrobiota, Proteobacteria (NB1-j) and Patescibacteria while suppressing Actinobacteriota and Firmicutes (Additional file 4). Regardless of the feedstock, 2.5% biochar influenced less and different microbial phyla than 0.25% biochar, generally increasing the abundance of Chloroflexi, Proteobacteria, Proteobacteria (NB1-j), and Sumerlaeota while decreasing the abundance of Firmicutes and the archaeal phylum Thermoplasmatota. At the class level, 0.25% biochar tended to enrich NB1-j (Proteobacteria phylum), OM190 (Planctomycetota phylum), and Vicinamibacteria (Acidobacteriota phylum), but suppressed Bacilli (Firmicutes phylum) and Actinobacteria (Actinobacteriota phylum). Biochar at 2.5% rate generally enriched Alphaproteobacteria and Gammaproteobacteria in the Proteobacteria phylum, while suppressing Bacilli (Firmicutes phylum), Holophagae (Acidobacteriota phylum), Microgenomatia (Patescibacteria phylum), and the archaeal class Thermoplasmata. Interestingly, wheat biochar enriched much more taxa, but did not inhibit more taxa, than other biochar types. Further, the same type of biochar affected different taxa at low vs. high rate.

The top 40 genera of the wheat rhizosphere accounted for 48.22% of all prokaryotes, with the top 20 genera being *Pseudomonas*, *Massilia*, *Sphingomonas*, *Bacillus*, *Pedosphaeraceae*, *Vicinamibacteraceae*, *Streptomyces*, *Allorhizobium-Neorhizobium-Pararhizobium-Rhizobium*, *Devosia*, *Arthrobacter*, *Altererythrobacter*, *Steroidobacter*, *Flavisolibacter*, *WD2101_soil_group*, *Lysobacter*, *Adhaeribacter*, *Blastococcus*, *Ohtaekwangia*, *Fluviicola*, *Longimicrobiaceae* (in descending order) (Fig S11), consistent with an earlier report of the rhizosphere core microbiome of cereal plants including wheat (66). Regardless of the feedstock and application rate, biochar consistently enriched *Ramlibacter* (Burkholderiales order) by 9.26−412.89%, *Longimicrobiaceae* (Longimicrobiales order) by 34.57−125.15%, *TRA3-20* (Burkholderiales order) by 16.26−77.93%, *Ellin6055* (Sphingomonadales order) by 10.19−57.35%, *Sphingomonas* (Sphingomonadales order) by 9.58−51.69%, and *Altererythrobacter* (Sphingomonadales order) by 0.98−43.68%, while inhibiting *Streptomyces* (Streptomycetales order) by 15.43−66.57%, *Arthrobacter* (Micrococcales order) by 42.99−73.85%, and *Bacillus* (Bacillales order) by 68.97−82.50% (Fig S12).

### Biochar enhanced microbial interactions in the wheat rhizosphere

Biochar treatments substantially enhanced microbial interactions, evident by the increased number of nodes, edges (except for C2.5, P2.5 and W2.5) and degree, larger connectivity, shorter path, smaller betweenness, and greater modularity (Table S2). These changes, i.e., small-world behavior, reflected strong synchronization among microbes in response to biochar-induced rhizosphere alterations. Further, positive correlations were predominant (>90%) in biochar-treated networks, suggesting mutualism and/or synergism due to corporations and functional dependency in the newly evolved rhizosphere.

Comparing the four biochar types, wheat biochar increased microbial network modularity more than other types and this effect was most significant for W0.25, likely reflecting the formation of specific functional modules (Table S2). Comparing the two application rates, 0.25% treatments (except for M0.25) resulted in greater increases in network degree than 2.5% treatments, suggesting higher microbial interactions under 0.25% than 2.5% biochar in general (Table S2).

Hub network analysis indicated that in the native wheat rhizosphere, 18 genera from six dominant phyla, as well as Cyanobacteria and Patescibacteria, had the highest betweenness centrality and thus were keystone taxa (Additional file 5) (Fig 5). New keystone taxa emerged across biochar treatments, including genera from four dominant phyla (Acidobacteriota, Myxococcota, Gemmatimonadota, Chloroflexi) and many phyla that comprised <5% of the rhizosphere microbiome (Armatimonadota, Bdellovibrionota, Deinococcota, Desulfobacterota, Elusimicrobiota, Fibrobacterota, Methylomirabilota, Sumerlaeota, WPS-2, WS2, Euryarchaeota, and Thermoplasmatota). This effect was most prominent in W0.25 treatment (29 keystone genera from 10 phyla), followed by M2.5 (27 keystone genera from 10 phyla). The newly emerged keystone taxa not only reflect community structure shifts but also suggest functional changes under specific treatments. For example, methylotroph *Methyloceanibacter* was keystone in C2.5 hub network; methanogen *Methanobacterium* (Euryarchaeota phylum) was keystone under M0.25 and M2.5 treatments; methylotroph *Methylotenera* (Proteobacteria) was keystone in P2.5 hub network. Similarly, nitrite-oxidizing bacterium *Nitrospira* was seen in W2.5 hub network, while sulfur (S)-oxidizing bacterium *Sulfurifustis* was seen in M0.25 and M2.5 hub networks. Further, networks evolved under biochar treatments typically had a large number of diverse keystone taxa with equally high centrality while the control hub network had two genera showing the highest centrality, indicating differential assembly of the rhizosphere microbiome due to biochar treatments.

**Fig 5.**
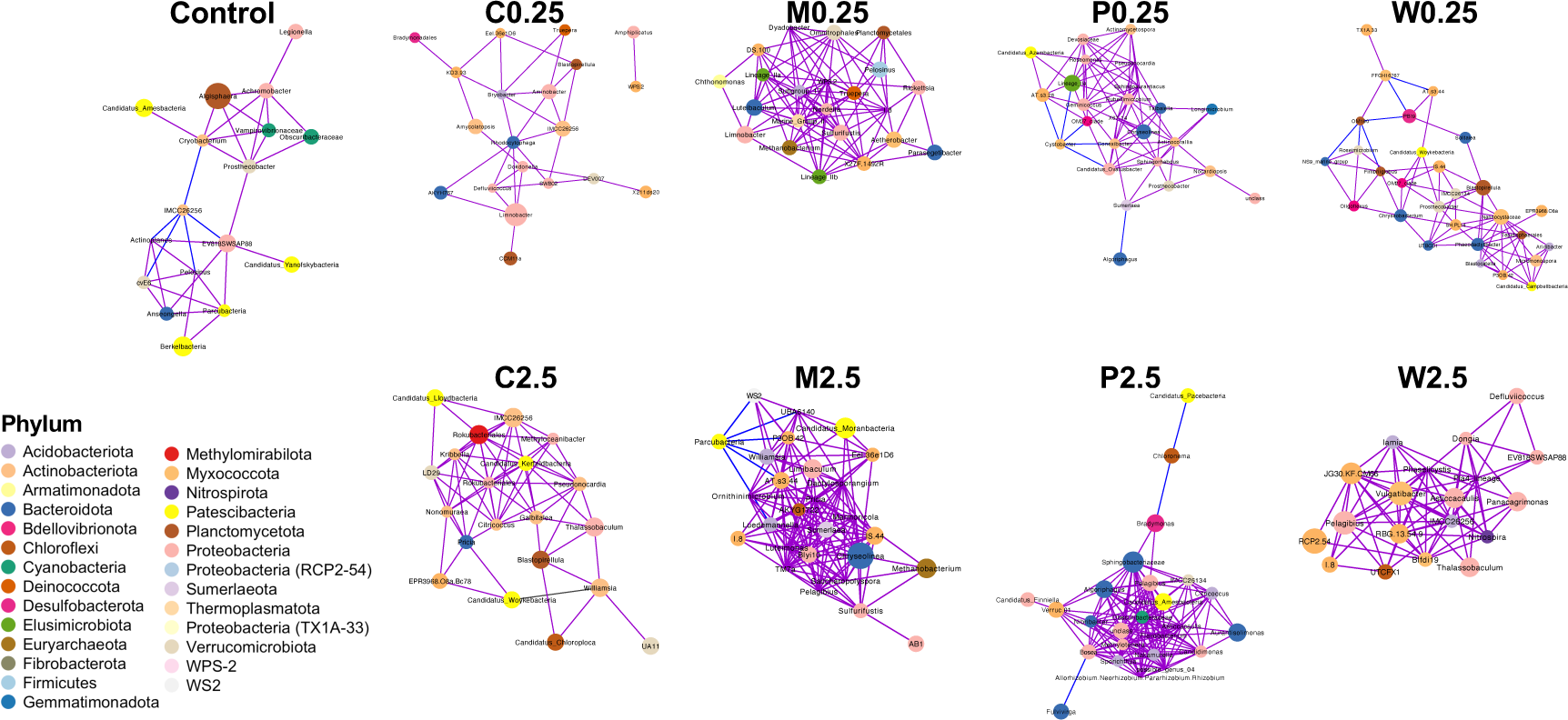
Hub network of microbial families/genera constructed using SparCC. Hub nodes were those above 95% quantile of the fitted log-normal distribution of three network centrality measures. Only significant correlations (*p* < 0.05) were retained here. Node color indicates the microbial phylum. Node size is proportional to the betweenness value. Purple and blue edges indicate positive and negative correlations, respectively.

### Biochar induced functional shifts in the rhizosphere microbiome

PICRUSt2 analysis showed biochar altered microbial functions in a treatment-dependent way, with W0.25 and M2.5 showing the strongest effects (Fig 6 and S13-S16, Additional file 6). W0.25 significantly influenced 48 microbial pathways, more than other treatments, and 32 pathways were unique changes to W0.25 (Fig 6A) (DESeq2: FDR-adjusted *p* < 0.01). M2.5 influenced 30 microbial pathways, and 17 pathways were unique changes to M2.5. Common changes seen in at least 3 different treatments included phosphotransferase system (PTS) and glycerolipid metabolism modulated by M0.25, P0.25, W0.25, and M2.5; flavone and flavonol biosynthesis modulated by C0.25, M0.25 and P2.5; flavonoid biosynthesis, and nucleocytoplasmic transport modulated by W0.25, C2.5, and M2.5; taurine and hypotaurine metabolism modulated by P0.25, C2.5, and M2.5; pyruvate metabolism, ascorbate and aldarate metabolism, and atrazine degradation modulated by M0.25, P0.25, and W0.25. Changes unique to W0.25 included upregulated translation, photosynthesis, biosynthesis of other secondary metabolites (flavonoids, isoflavonoids, indole alkaloids), biosynthesis of terpenoids and polyketides (vancomycin group antibiotics, zeatin), and glycan biosynthesis and metabolism, as well as downregulated amino acid metabolism, tropane, piperidine and pyridine alkaloid biosynthesis, fatty acid degradation, limonene degradation, degradation of xenobiotics (e.g., polycyclic aromatic hydrocarbons, chloroalkane/chloroalkene, caprolactam), among others (Fig S13-S16, Additional file 6). Changes unique to M2.5 included upregulated replication and repair, phosphatidylinositol signaling system, and carotenoid biosynthesis, as well as downregulated biosynthesis of other secondary metabolites (penicillins and cephalosporins, isoquinoline alkaloids, streptomycins), lipid metabolism, among others. W0.25 frequently showed contrasting effects than other treatments: W0.25 upregulated the abundance of nucleocytoplasmic transport and flavonoid biosynthesis pathway while others down-regulated the two pathways (DESeq2: FDR-adjusted *p* < 0.01) (Fig 6B, Additional file 6).

**Fig 6.**
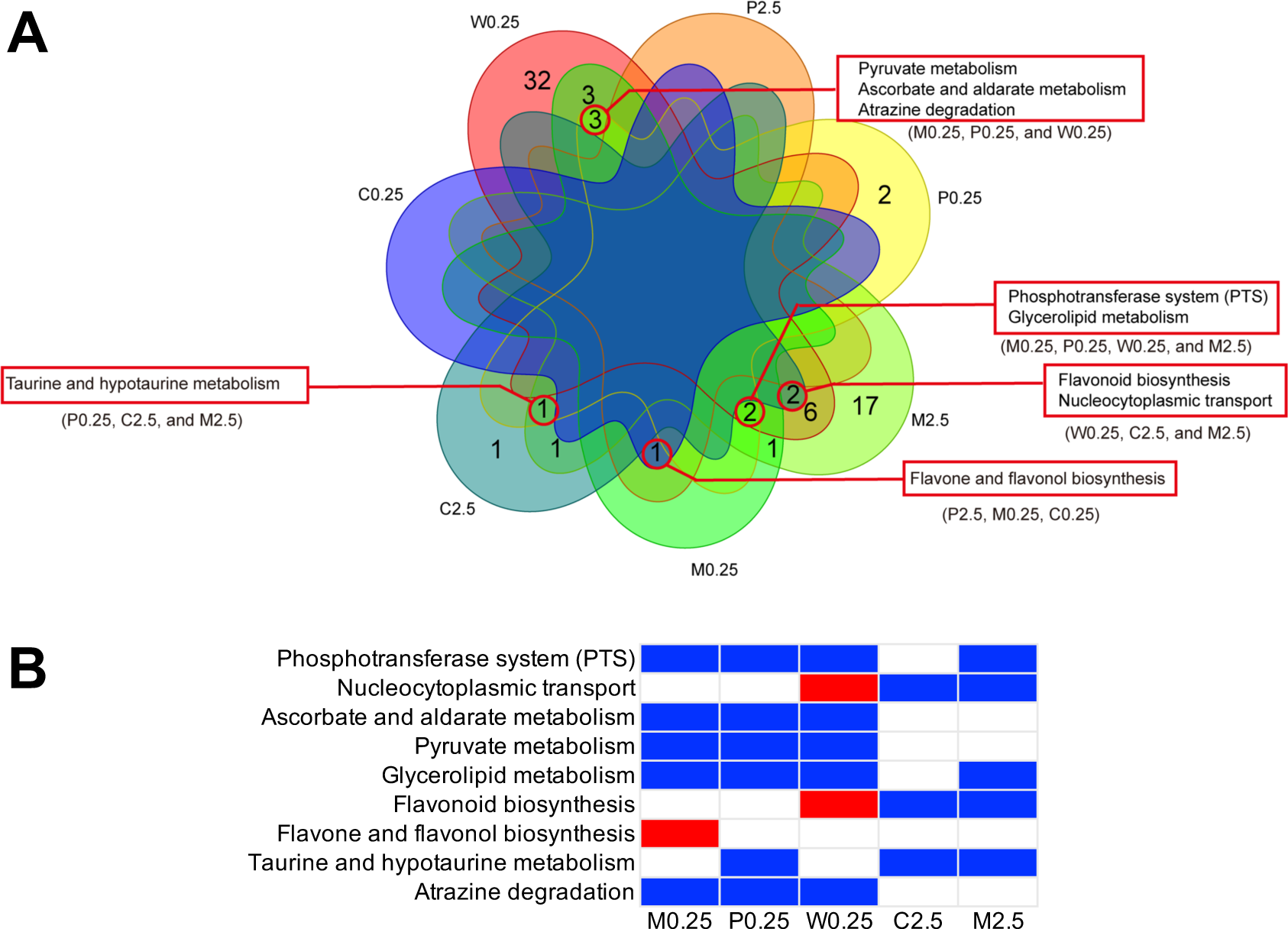
Microbial pathways significantly regulated by biochar treatments. W2.5 resulted in no significant changes and are not shown. (A) Venn diagram of common and specific alterations. (B) W0.25 strongly influenced more microbial pathways and often showed distinct effects compared to other treatments. Heatmap shows pathway alterations identified in 3-4 out of 7 different treatments. Red, upregulation; blue, downregulation.

Given the strong and distinct effects of W0.25 compared to other treatments (Fig 6 and S13-S16, Additional file 6), we used multi-omics network analysis to explore potential correlations between root metabolites and rhizosphere microbial genera whose functions may be involved in processing those metabolites. Eighteen significantly altered microbial pathways and the associated genera were used to construct microbe-metabolite co-occurrence networks (Fig 7) (Additional file 7). Compared to the control network where *Pseudomonas* had higher centrality than *Bacillus*, the W0.25 network showed similar centrality for these two genera. Connectivity between other less dominant genera and root metabolites also increased in the W0.25 network compared to the control network.

**Fig 7.**
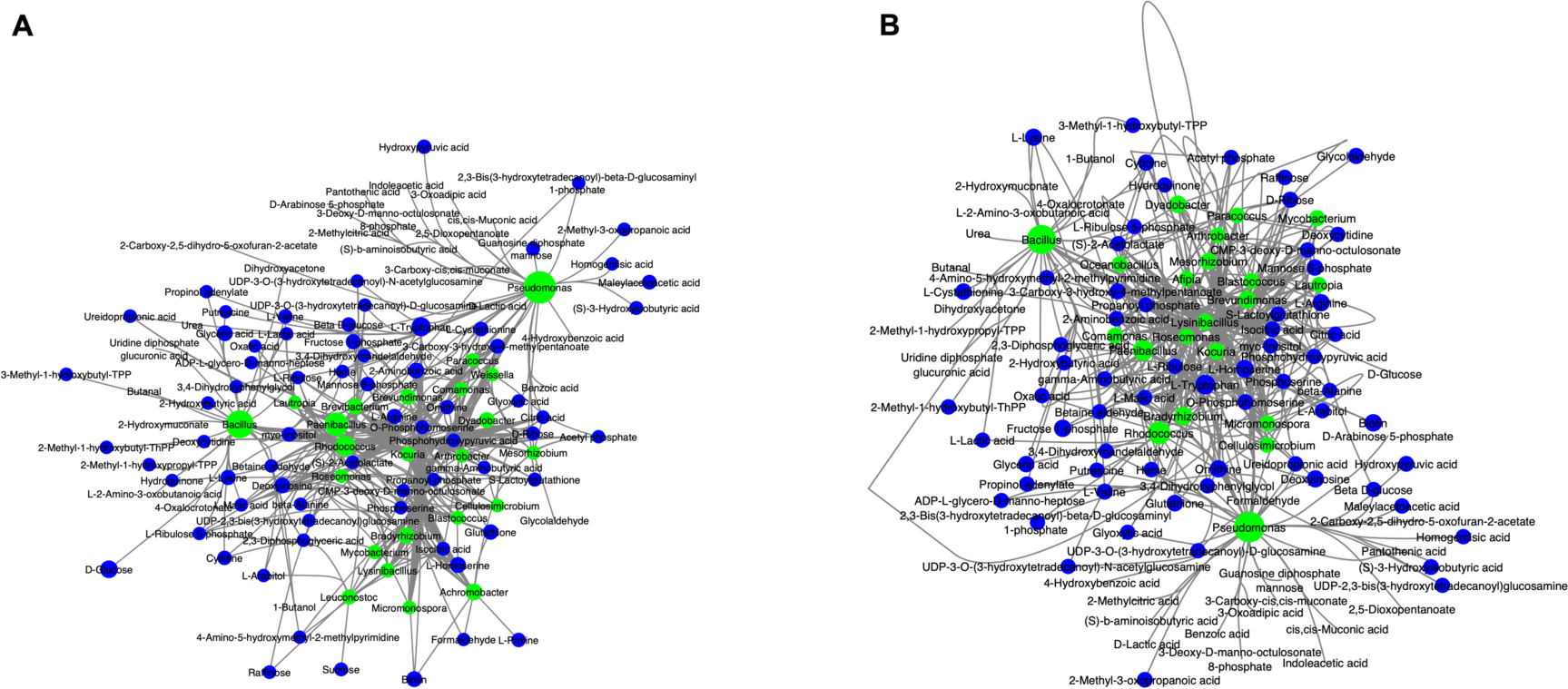
Microbial genera-root metabolite co-occurrence network under control (A) and W0.25 (B). Only significant microbial pathways (FDR-adjusted *p* < 0.01 in both OmicsNet 2.0 and PICRUSt2) were used for network construction. Green nodes indicate microbial genera; blue nodes indicate root metabolites. Node size is proportional to the betweenness value.

### Influential factors of biochar’s modulating effects

To understand why biochar modulated various aspects of the rhizosphere in a feedstock-dependent way, we analyzed biochar surface element composition and the content of three oxygen-containing functional groups (acidic carboxyl, lactone, and phenolic group). Carbon (C%) and oxygen (O%) atoms accounted for ∼90% of all elements on the biochar surface (Additional file 8). Manure biochar had a significantly higher amount of all three oxygen-containing functional groups than other types of biochar (Fig S17) (ANOVA: *p* < 0.05). Corn, pine, and wheat biochar had similar contents of lactone and phenolic groups. Pine and corn biochar showed the highest and lowest acidic carboxyl content, respectively. Surface C% and O% and the three oxygen-containing functional groups partially explained the observed variations in root metabolites (NMDS: stress ∼ 0.18), and surface C% significantly influenced root metabolites, particularly non-glycans (PERMANOVA: *p* < 0.005) (Fig S18A-C). Biochar surface properties also influenced the rhizosphere microbiome (NMDS: stress = 0.087), and O% and oxygen-containing groups significantly influenced the microbiome composition (PERMANOVA: *p* < 0.02) (Fig S18D).

## Discussion

### Biochar modulated wheat root metabolism and likely the rhizosphere chemistry

Biochar is a carbonaceous material typically produced by pyrolysis of various biomass under oxygen-limiting conditions. Biochar soil amendment has the potential to simultaneously sequester C and provide multiple benefits to the soil-plant continuum (70). Yet, the underlying mechanisms are not fully understood and how biochar modulates the plant rhizosphere remains elusive (71). Using wheat as a model plant, we revealed biochar’s profound effect in reshaping the plant rhizosphere.

Biochar modulated a broad range of metabolic pathways in wheat roots, which likely changed the chemical gradient of the rhizosphere. Twenty-five pathways were regulated, including 6 pathways in amino acid metabolism, 5 pathways in biosynthesis of other secondary metabolites, 4 pathways in metabolism of cofactors and vitamins, 4 pathways in metabolism of terpenoids and polyketides, 2 pathways in lipid metabolism, and 1 pathway in each of carbohydrate metabolism, glycan biosynthesis and metabolism, nucleotide metabolism, and translation (Fig 1-3 and S2) (Additional file 1-3). Changes in amino acid metabolism, carbohydrate metabolism, pyrimidine metabolism, and biosynthesis of secondary metabolites have also been identified in biochar-treated plants with enhanced resistance to abiotic stresses (72-75).

Using untargeted metabolomics, we revealed common changes induced by four types of biochar (corn, manure, pine, and wheat) and two application rates (0.25% and 2.5%), which provides a molecular basis for the widely recognized biochar effects to the soil-plant continuum. Amino acid metabolism (e.g., phenylalanine, tyrosine and tryptophan biosynthesis) was most commonly altered by biochar. Amino acids are building blocks of proteins and contribute to the biosynthesis of an enormous variety of secondary metabolites that are essential for plant growth and development, signaling, and defense against various stresses (76). In particular, phenylalanine, tyrosine, and tryptophan are involved in the production of a large number of secondary metabolites (e.g., glucosinolates, alkaloids, phenolics, terpenes) (77, 78). Alterations in amino acid metabolism could have cascade effects in the dynamics of secondary metabolites. Indeed, biosynthesis of other secondary metabolites was the second most commonly altered in this study, followed by metabolism of terpenoids and polyketides. Metabolism of cofactors and vitamins (e.g., riboflavin metabolism, ubiquinone and other terpenoid-quinone biosynthesis) was also commonly altered. Regulation of the aforementioned pathways seems a common mechanism for wheat to respond to abiotic and biotic stimuli, and stress resistance of certain wheat cultivars is associated with differential expression of these pathways (79, 80).

A wide range of plant signaling metabolites from the category of other secondary metabolites and terpenoids and polyketides were regulated by all biochar treatments, including phenylpropanoids, flavonoid compounds, anthocyanins, glucosinolates, brassinosteroids, carotenoids, diterpenoids, and sesquiterpenoids and triterpenoids (Additional file 2). These compounds are involved in plant-environment interactions and used by plants to inhibit pathogens or recruit plant growth-promoting bacteria (PGPB) (77, 81). For example, flavonoids are phenolic secondary metabolites derived from phenylalanine with a wide range of roles in plant development, defense, and interaction with other organisms including microbes (82, 83). Flavonoids broadly contribute to the rhizosphere microbiome diversity and preferentially attract plant-specific PGPB, resulting in bacteria-enhanced plant resistance to adverse conditions and fungal pathogens (84-86). Brassinosteroids are steroidal plant hormones functioning as a master regulator, able to regulate root growth through localized biosynthesis along the longitudinal root axis and modulate plant interactions with pathogenic microbes (81, 87). Besides, all biochar treatments regulated metabolism of cofactors and vitamins, including ubiquinone and other terpenoid-quinone biosynthesis, that can also be involved in plant-PGPB interactions (88). Further, ubiquinone functions as an electron transporter on the mitochondrial membrane during aerobic respiration and the catabolism of certain branched-chain amino acids. It is also involved in plant response to stress by scavenging reactive oxygen species (ROS), and may indirectly regulate cell signaling and gene expression by mediating the generation of hydrogen peroxide, an important signaling molecule in plant resistance and cell metabolism (88, 89). In line with these common pathway alterations, metabolite networks had substantially increased connectivity after all biochar treatments, reflecting profound effects of altered amino acid metabolism to the production of secondary metabolites. Taking together, biochar soil amendment induced a systemic response and complex regulation in the plant root which could directly influence rhizosphere chemistry. Future research should apply plant transcriptomics and proteomics to further attest this connection. Nevertheless, these common metabolic alterations may explain biochar’s multiple benefits to plant productivity, including root system development, and some of them have also been observed in PGPB-treated wheat (e.g., phenylalanine metabolism, flavone and flavonol biosynthesis, riboflavin metabolism, ubiquinone and other terpenoid-quinone biosynthesis) (70, 90, 91).

### Biochar reshaped the wheat rhizosphere microbiome

Numerous studies have reported biochar-induced microbiome changes in bulk soil, as discussed in our recent systematic literature review and meta-analysis, but how biochar modulates the rhizosphere is still obscure (8). Our results indicate that regardless of feedstock and application rate, biochar increased rhizosphere microbial diversity (Fig S3-S8), altered microbiome composition (Fig 4 and S9-S12, Additional file 4), enhanced microbe-microbe interactions (Fig 5), and resulted in functional changes. We speculate these changes were due to complex interactions between plant roots and soil microbes upon biochar amendments. This speculation is supported by our metabolomics data as many compounds in the biochar-altered pathways are major plant exudates, and some of them, including amino acids, various secondary metabolites, and vitamins, are involved in plant-microbe bidirectional interactions (76, 83, 88). For example, auxotrophy for specific amino acids is a selective fitness advantage for certain plant-beneficial rhizosphere microbes (14). Flavonoids and their degradation products have multifaceted roles in mediating the assembly and functioning of rhizosphere microbiomes, and are used by plants to recruit PGPB to enhance stress tolerance (83). Vitamins like riboflavin and degradation products have multiple roles in plant-microbe interactions, including the activation of bacterial quorum-sensing receptors (88).

Increased microbial diversity is a common phenomenon in the rhizosphere after stimulation by carbonaceous nanomaterials (28), and this typically leads to divergent microbiomes depending on the stimulus material, which was also observed here (Fig 4 and S9). Biochar treatments frequently enriched Elusimicrobiota and Proteobacteria (particularly Alphaproteobacteria, Gammaproteobacteria and the Deltaproteobacteria order NB1-j) and genera in the order Burkholderiales and Sphingomonadales, while suppressing Firmicutes (particularly Bacillales order) and Thermoplasmatota (Thermoplasmata class), as well as the genus *Streptomyces* and *Arthrobacter* (Fig 4 and S10-S12, Additional file 4). These changes reflect the adaptation of the rhizosphere microbiome to a biochar-altered microenvironment, which could lead to functional shifts. Evidence from metagenome-assembled genomes suggests Elusimicrobiota may participate in glycan biosynthesis and metabolism, carbohydrate metabolism, as well as biosynthesis of essential amino acids, B vitamins, and niacin and nicotinamide (92). NB1-j may participate in N fixation (*nifD*), nitrification (*hao*), and denitrification (*norB*) (93), while Burkholderiales could contribute to complete denitrification, particularly reduction of N_2_O to N_2_ catalyzed by the NOSZ reductase (94). These results are consistent with common reports that biochar could increase nitrification and yet still reduce N_2_O emission (8). Further, Burkholderiales can be free-living in the rhizosphere or endophytic and consists of PGPB species able to enhance nodulation and N fixation, sequestrate iron and mobilize inorganic phosphate using siderophores, compete against plant pathogens by secreting antibiotics, produce phytohormones, and trigger induced systemic resistance (95). A recent study by Chen et al. (2019) found the abundance of Burkholderiales in the wheat rhizosphere correlated positively with organic acids in root exudates, which themselves varied with wheat growth stage and N fertilization level (11). Another study showed that elevated CO_2_ and nitrate level jointly influenced genes related to denitrification, amino sugar and nucleotide sugar metabolism, fructose and mannose metabolism, pyruvate metabolism, lipid biosynthesis, biofilm formation, and secretion systems in the wheat root endosphere microbiome, and that Burkholderiales were responsible for these functional changes. Our results support the previous studies and suggest Burkholderiales responded to biochar-induced changes in rhizosphere chemistry and thereby influenced N-cycling therein. Our observations of a general increase in Alphaproteobacteria, Gammaproteobacteria, and Sphingomonadales and a decrease in *Streptomyces* in the ectorhizosphere of tillering stage wheat (i.e., 15 days here) after biochar amendment are also consistent with those seen in wheat with increasing N fertilization (96). However, the same study found increasing N fertilization stimulated *Arthrobacter* in the rhizosphere and root endosphere, as well as on root surface, of wheat (96), opposite to the trend of *Arthrobacter* observed in this study. Therefore, our observations were not solely due to biochar-introduced changes in soil N. Indeed, here biochar reduced the Bacillales order, while Bacillales increased after intensified N fertilization (96). Biochar suppressed archaeal methanogens (Thermoplasmata class), in line with widely reported mitigation of methane emissions by biochar (97). This was likely due to biochar’s liming effect (8), which inhibited the acidophilic Thermoplasmata class (98).

### Biochar-induced rhizosphere microbiome assemblage and functional shifts therein

Remarkably, biochar increased microbial network complexity, especially modularity, in the wheat ectorhizosphere (Fig 5, Additional file 5), reflecting biochar-induced microbiome assembly and the formation of specific functional modules (99, 100). It is generally believed that system stability increases with complexity, particularly modularity, and that higher complexity and linked functions make a community less vulnerable to environmental changes (101). Increased network complexity was previously seen in biochar-treated bulk soil (102, 103). In contrast, microbiome assemblage in the ectorhizosphere, the narrow interface between the outermost region of roots and the bulk soil, is strongly influenced by root exudates. Our findings of rhizosphere microbiome and root metabolism align with this common knowledge, and contribute to the still limited literature on biochar-manipulated plant-microbe interactions. Further, biochar increased positive correlations among microbes, suggesting strong microbial synergism which allows a microbiome to quickly adapt to a new environment (100). On the contrary, increased ratios of negative to positive correlations between taxa were seen in microbiomes of biochar-treated tobacco (103) and those from decreasing environmental stresses (104). Here, biochar’s stronger effects on positive correlations could result from habit filtering (105). Together, our results suggest biochar fostered a more stable rhizosphere microbiome, with many phenomena similar to the effects of long-term organic amendments (106-108), and thus promote plant growth and health.

Hub network analysis has the potential to identify keystone taxa (109). We found in the control hub network, two genera were the dominant keystone taxa and one of them (Pelosinus) likely mediated iron reduction (Pelosinus) (Additional file 5) (110). In contrast, biochar treatment hub networks had a larger number of diverse keystone taxa from both dominant and non-dominant phyla, and many of them had similar degree, closeness, and betweenness values. This suggests a decentralized rhizosphere microbiome and a transition to a small-world network consisting of specific modules(100), which is supported by the presence of keystone taxa involved in methane, N, and S cycling. These phenomena are similar to those observed for organic amendments, where soil microbiomes were decentralized into small modules of functionally interrelated taxa(106). Interestingly, archaea were rarely keystone taxa in this study, except for mature biochar, consistent with an earlier report that neutral processes rather than deterministic factors drive soil archaeal communities (111).

### Influential factors of biochar’s rhizosphere modulating effects

Both application rate and feedstock choice strongly influenced biochar’s rhizosphere modulating effects. For all four types of biochar, a lower rate (0.25%) enhanced wheat growth, whereas a higher rate (2.5%) inhibited growth, consistent with other reports of rate-dependent biochar effects on plant physiology, including the development of root systems (112). Additionally, our study provides new insights into the underlying mechanisms: 2.5% biochar commonly affected 71 root metabolic pathways, while 0.25% biochar commonly affected 29 pathways; however, 2.5% biochar influenced less and different microbial phyla than 0.25% biochar (Additional file 2-5). Treatment-dependent root metabolic differences are largely reflected in the regulation of 44 metabolites from 13 classes (Fig 2), predominantly consisting of carboxylic acids and derivatives, flavonoids, organooxygen compounds, and prenol lipids that are known to be involved in plant root-microbe interactions (83, 113, 114). Biochar type exerted greater influence than application rate (Fig 1, S2 and S18), and some treatments showed unique effects, such as glycosaminoglycan degradation regulated only by W0.25.

Coincidently, the rhizosphere microbiome also showed feedstock-specific alterations, both in structure (Fig 4 and S9-S12) and functioning (Fig 6 and S13-S16). Many of the predicted functional changes are consistent with the observed root metabolic alterations, including carbohydrate metabolism, the PTS, biosynthesis flavonoid compounds, among others. The PTS is a carbohydrate transport system and also regulates numerous cellular processes, including carbon catabolite repression, stress response, and biofilm formation (115). Regulation of the PTS and carbohydrate metabolism could reflect adaptation of the rhizosphere microbiome to biochar-induced carbohydrates in root exudates. Flavonoid compounds can modulate the dynamic structure and functioning of the rhizosphere microbiome, determine patterns of root growth, and broadly affect plant growth, development, and defense. *Pseudomonas* and *Rhizobium*, two dominant genera detected in this study, could actively degrade flavonoid molecules, thus mediating the dynamics of flavonoid compounds in the rhizosphere (83). Future research should use soil metagenomics and metatranscriptomics to validate our results, and empirical evidence is needed to directly prove biochar-induced, root exudates-influenced rhizosphere microbiome assembly and the formation of specific functional modules therein.

Remarkably, W0.25 had strongest and distinct modulating effects on both root metabolism and rhizosphere microbiome, and enhanced root metabolite network connectivity and microbial network modularity more than other treatments (Fig 3 and 5, Table S1-S2). This suggests the induction of orchestrated chemical and microbial changes in the rhizosphere. Many of W0.25-unique microbial functional alterations are relevant to plant-microbe interactions, including upregulation of flavonoid biosynthesis, isoflavonoid biosynthesis, indole alkaloid biosynthesis, and zeatin biosynthesis, as well as downregulation of limonene degradation (Fig 6 and S16, Additional file 6). Accumulation of flavonoids have been seen in wheat cultivars resistant to fungal diseases and PGPB-treated wheat plants with priming defense (90, 116, 117). The same mechanism may have occurred to W0.25-treated wheat, where the newly assembled rhizosphere microbiome contributed to the dynamics of flavonoids at the plant-microbe interface. Upregulated indole alkaloid biosynthesis could be a result of metabolism of tryptophan in root exudates by *Bacillus* spp., which may in turn contribute to enhanced resistance to insects in wheat (118, 119). Similarly, rhizosphere microbes like *Bacillus subtilis* can metabolize tryptophan and other small molecules in root exudates and produce the zeatin cytokinin, a phytohormone, thus stimulating plant growth and further amino acid deposition by wheat roots (120). Limonene is a volatile monoterpenoid with antimicrobial properties (121), and exposure to limonene can suppress fungal disease (122). Thus, downregulated limonene degradation could protect wheat from disease. This change may have occurred through W0.25 decreasing *Rhodococcus* spp. (Actinobacteria class) that possess limonene degradation pathways (123). Consistently, W0.25 suppressed the Actinobacteria class. More evidence of W0.25-mediated plant-microbe interactions came from multi-omics network analysis, which showed increased connectivity between root metabolites and microbial genera with relevant functions (18 pathways) after W0.25 treatment (Fig 7, Additional file 7). Elucidating the detailed chemical interactions between roots and microbes remains challenging, and future research should integrate soil metagenomics and metabolomics to explore microbial metabolism of biochar-induced root exudates. Regardless of the exact mechanisms, feedstock-dependent biochar surface chemistry (i.e., C%, O%, acidic carboxyl, lactone, and phenolic group) partially drove differential alterations in the rhizosphere (Fig S18), which has implications for targeted biochar application in sustainable agriculture. Elucidating the detailed chemical interactions between roots and microbes remains challenging, and future research should integrate soil metagenomics and metabolomics to explore microbial metabolism of biochar-induced root exudates. Regardless of the exact mechanisms, feedstock-dependent biochar surface chemistry (i.e., C%, O%, acidic carboxyl, lactone, and phenolic group) partially drove differential alterations in the rhizosphere (Fig S18), which has implications for targeted biochar application in sustainable agriculture.

### The potential of carbonaceous material-based rhizosphere microbiome engineering

We revealed how four types of biochar, applied at two rates, modulated the wheat rhizosphere by inducing common and unique changes in root metabolites and microbiome structure and functioning. These findings contribute to our mechanistic understanding of frequently reported benefits of biochar amendment to the soil-plant continuum (8, 70). Our observations of orchestrated chemical and microbial changes in the rhizosphere provide new insights into the potential of rhizosphere microbiome engineering using biochar or other carbonaceous materials (18-20). The rhizosphere is a dynamic microenvironment shaped by entangled webs of interactions between the plant and its surrounding soil. Different types of biochar induced differential systemic responses of the plant-microbiome holobiont. Induced root metabolites reshaped the assembly of the rhizosphere microbiome by serving as chemoattractants or substrates for microbes (76). The new assemblage constituted microbes better adapted to biochar-modulated rhizosphere, including those capable of utilizing biochar-induced root metabolites, and had a highly modular network structure where keystone taxa played essential roles in metabolism of nutrients or root metabolites (100). The reprogramed rhizosphere microbiome functioning could in turn benefit the host plant by promoting nutrient acquisition and stress tolerance (121). Therefore, biochar amendment represents a top-down *in situ* manipulation approach that elicits a transition of the native microbiome toward optimal survival mode through ecological selection (124). Biochar holds the promise of engineering the rhizosphere microbiome by reprogramming root-microbe interactions—understanding these relationships between biochar physicochemistry and beneficial root-microbe interactions will ensure the realization of this promise towards a sustainable agriculture.

### Conclusions

Using wheat as a model plant, we revealed that biochar could modulate plant root metabolites and the rhizosphere microbiome. Comparisons of four types of biochar and two rates identified common and unique modulating effects of these treatments. Regulation of amino acid metabolism was the most common effect on root metabolism, which had cascade effects in the dynamics of a wide range of secondary metabolites, including many plant signaling molecules (i.e., other secondary metabolites, terpenoids, and polyketides) that are known to be involved in plant-environment interactions and used by plants to inhibit pathogens or recruit PGPB. All biochar treatments increased rhizosphere microbial diversity, altered community composition, enhanced microbial interactions, and resulted in functional changes. Increased Burkholderiales (denitrifying bacteria) abundance and decreased Thermoplasmata (archaeal methanogens) abundance could explain biochar’s widely reported effects on N_2_O and CH_4_ mitigation, respectively. Biochar enhanced positive correlations among microbes and network complexity, suggesting local adaptation through mutualism and/or synergism and the formation of modules of functionally interrelated taxa. A large number of diverse keystone taxa from both dominant and non-dominant phyla emerged after biochar treatments, including many known to be involved in CH_4_, N, and S cycling. Treatment-specific alterations also occurred, and biochar feedstock exerted greater influence than application rate. W0.25 showed the strongest and most distinct modulating effects, resulting in orchestrated changes in both root metabolites and rhizosphere microbiome, especially those relevant to plant-microbe interactions and likely beneficial to the host plant. Our work contributes to a mechanistic understanding of how biochar modulates the soil-plant continuum. Future research utilizing integrated multi-omics is needed to directly demonstrate root exudate-microbe interactions. Still, this work provides new insights into the potential of top-down rhizosphere microbiome engineering through carbonaceous material-based reprogramming of root-microbe interactions.

## Declarations

### Ethics approval and consent to participate

Not applicable.

### Consent for publication

Not applicable.

### Availability of data and materials

All Illumina reads have been deposited at the NCBI Sequence Read Archive (SRA), under the BioProject accession number PRJNA1136486.

### Competing interests

The authors declare that they have no competing interests.

### Funding

This work was funded by Idaho State University through a Developing Collaborative Partnerships grant, the State University of New York College of Environmental Science and Forestry, and the US Department of Agriculture through a McIntire-Stennis Capacity Grant (NI23MSCFRXXXG070). Y. You also received support from the Center for Advanced Energy Studies (CAES), a research, education, and innovation consortium consisting of Idaho National Laboratory and the public research universities of Idaho. H. Yang is grateful to the ESF Alumni Association. P. Kerner is grateful to Idaho State University Center for Ecological Research and Education.

### Authors’ contributions

YY conceived the project and obtained funding. ES and AM produced biochar. HY conducted biochar surface characterization. XL and YY designed the pot experiments, and XL conducted the experiments. XL and YY processed the plants and collected the data. PK processed the rhizosphere soils, extracted DNA, and assisted in Illumina sequencing. PK, HY, and YY performed bioinformatics. HY and YY performed data analysis and wrote the manuscript. All authors read, revised and approved the submitted version of the manuscript.

## Acknowledgements

We thank Dr. Yaqiao Wu at the Microscopy and Characterization Suite (MaCS) of the Center for Advanced Energy Studies (CAES) for assisting in biochar characterization, Dr. Xinzhi Pu at the Biomolecular Research Center of Boise State University for assisting in LS-MS analysis, and Lisa McDougall and Jason Werth at the Molecular Research Core Facility of Idaho State University) for completing Illumina sequencing. We are grateful to Syracuse University Research Computing for providing access to a high-performance computing cluster.

## Notes

### Competing Interest Statement

The authors have declared no competing interest.

