## Supplemental Information for "Biochar Modulates Wheat Root Metabolome and Rhizosphere Microbiome in a Feedstock-dependent Manner"

Number of figures: 20

### Supplementary Methods

#### Biochar characterization

Three oxygen-containing functional groups (acidic carboxyl, lactone, phenolic group) were determined by Boehm Titration (1, 2). Briefly, 0.2 g of a biochar sample was added to 20 mL of 0.05 M $NaHCO_3$, 0.1 M $Na_2CO_3$, or 0.1 M NaOH, respectively. The mixture, along with a no-biochar blank control of the same solution, was shaken for 24 h and then filtered through a Whatman No. 42 filter paper to remove solid particles. Subsequently, an aliquot (10 mL) of the filtrate was mixed with 15 mL of 0.1 M HCl to ensure complete neutralization of the bases. The resulting solution was then back-titrated with 0.1 M NaOH solution, using phenolphthalein as an indicator. The total surface acidity was determined by calculating the moles neutralized by NaOH. The carboxyl group content was determined by the moles neutralized by $NaHCO_3$. The lactone group content was calculated as the difference between the values obtained from the $NaHCO_3$ and $Na_2CO_3$ titrations. The difference between molar NaOH and $Na_2CO_3$ was used to represent the phenolic group content, as described by Rutherford *et al.* (2008) (3).

#### Metabolic data processing and analysis

When integrating the compound tables obtained from different platforms into a single table, in cases where a compound was detected by multiple platforms, we used the peak value assigned by XCMS for that compound. This is because typically, XCMS yielded the highest recognition peak and identified the widest range of compound types among the four platforms used in this study. If a compound was not recognized by XCMS but identified by MetaboAnalyst 5.0 or other platforms, we use the recognition peak value from MetaboAnalyst 5.0 because this could facilitate downstream functional enrichment analysis in MetaboAnalyst 5.0.

### Supplementary Tables

**Table S1.** Topological properties of root metabolite co-occurrence networks under different treatments. Values reflect averages across biological replicates.

| Treatment | Nodes | Edges | Average degree | Degree centrality | Connectivity |
|---|---|---|---|---|---|
| Control | 424 | 601 | 2.8349 | 0.02167 | 0.0067 |
| C0.25 | 345 | 504 | 2.9217 | 0.0380 | 0.0085 |
| C2.5 | 365 | 411 | 2.2521 | 0.0130 | 0.0062 |
| M0.25 | 408 | 609 | 2.9853 | 0.0246 | 0.00733 |
| M2.5 | 379 | 580 | 3.0607 | 0.0263 | 0.0081 |
| P0.25 | 352 | 404 | 2.2955 | 0.0163 | 0.0065 |
| P2.5 | 345 | 490 | 2.8406 | 0.0266 | 0.0083 |
| W0.25 | 400 | 882 | 4.4100 | 0.0616 | 0.0110 |
| W2.5 | 372 | 551 | 2.9624 | 0.0298 | 0.0080 |

**Table S2.** Topological properties of microbial co-occurrence networks based on genus under different treatments. Values reflect averages across biological replicates.

| Treatment | Nodes | Edges | Average degree | Average path | Average connectivity | Average betweenness | Average clustering | Modularity |
|---|---|---|---|---|---|---|---|---|
| Control | 3960 | 432 | 18.33 | 2.73 | 0.0425 | 0.1185 | 0.3918 | 0.4373 |
| C0.25 | 6822 | 452 | 30.19 | 2.22 | 0.0669 | 0.0486 | 0.4471 | 0.4510 |
| C2.5 | 6110 | 426 | 28.69 | 2.38 | 0.0675 | 0.0314 | 0.4742 | 0.4398 |
| M0.25 | 6066 | 474 | 25.59 | 2.42 | 0.0541 | 0.0866 | 0.4055 | 0.4799 |
| M2.5 | 6950 | 461 | 30.15 | 2.35 | 0.0656 | 0.0549 | 0.4597 | 0.4449 |
| P0.25 | 6929 | 456 | 30.39 | 2.28 | 0.0668 | 0.0328 | 0.4349 | 0.4655 |
| P2.5 | 4894 | 429 | 22.82 | 2.51 | 0.0533 | 0.0498 | 0.4768 | 0.4825 |
| W0.25 | 6592 | 443 | 29.76 | 2.66 | 0.0673 | 0.0338 | 0.4167 | 0.5177 |
| W2.5 | 6114 | 422 | 28.98 | 2.68 | 0.0688 | 0.0479 | 0.4474 | 0.4937 |

### Supplementary Figures

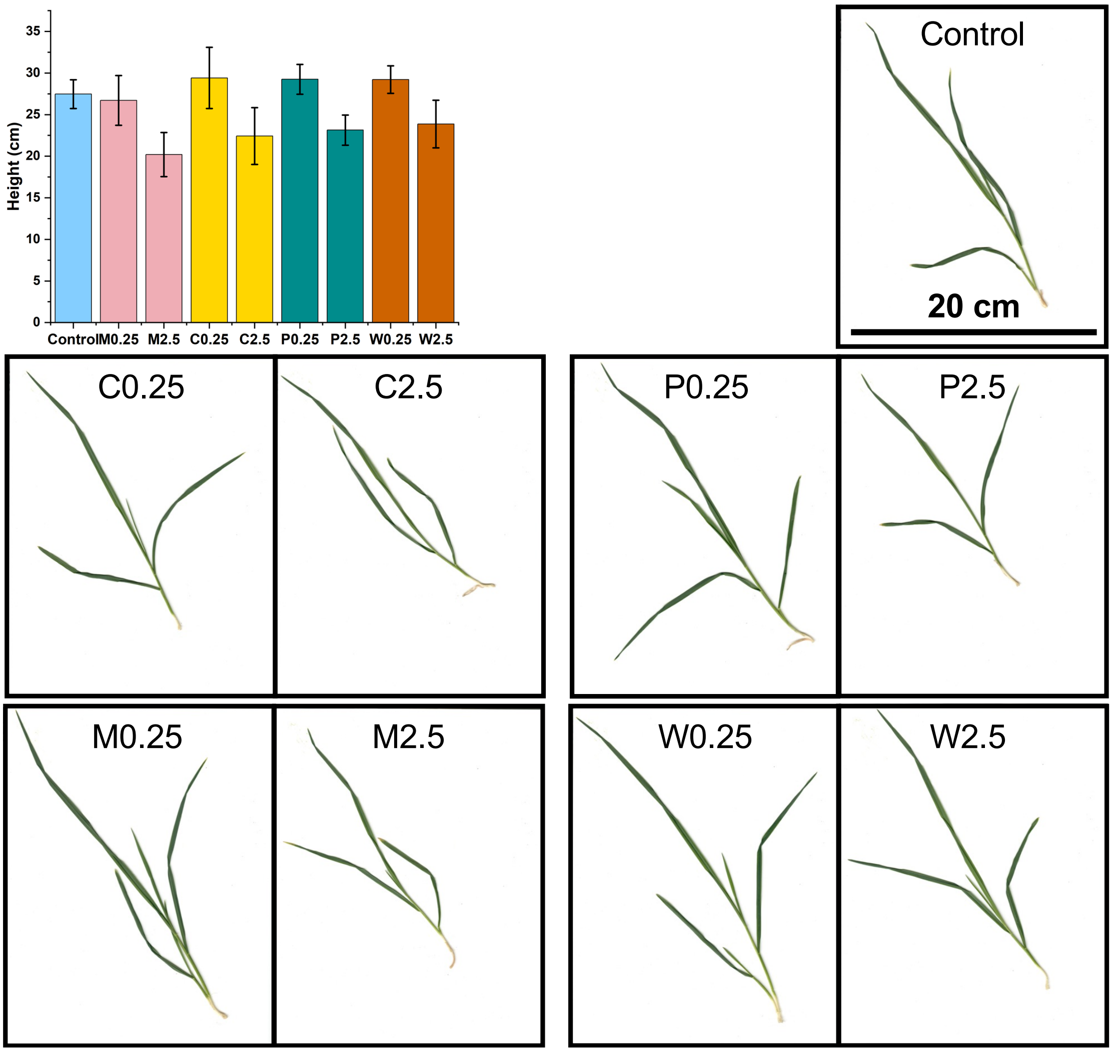


**Fig S1.** Height and shoot biomass of wheat plants harvested after 15-day pot experiments. Biochar treatments are indicated by feedstock (C, corn stover; M, cattle manure; P, pine sawdust; W, wheat straw) and application rate (0.25% or 2.5%).

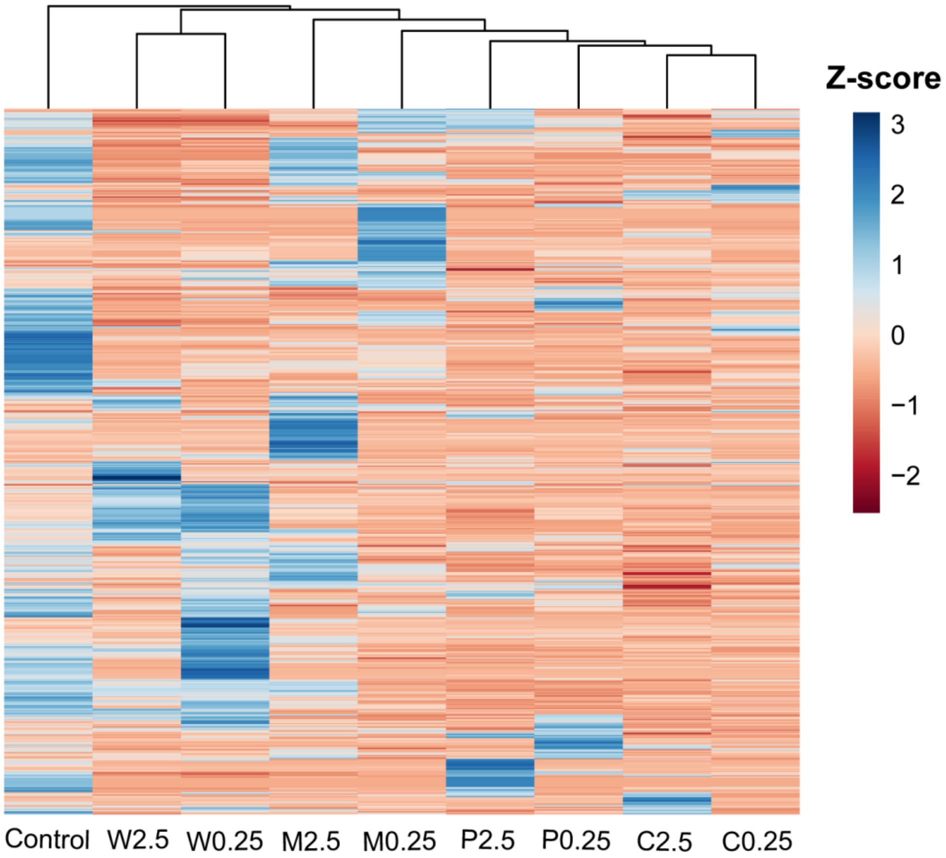


**Fig S2.** Heatmap of root metabolites under various treatments. Each row represents a metabolite. Blue and red colors indicate normalized MS signal intensities. Clustering is based on hierarchical method using complete linkage.

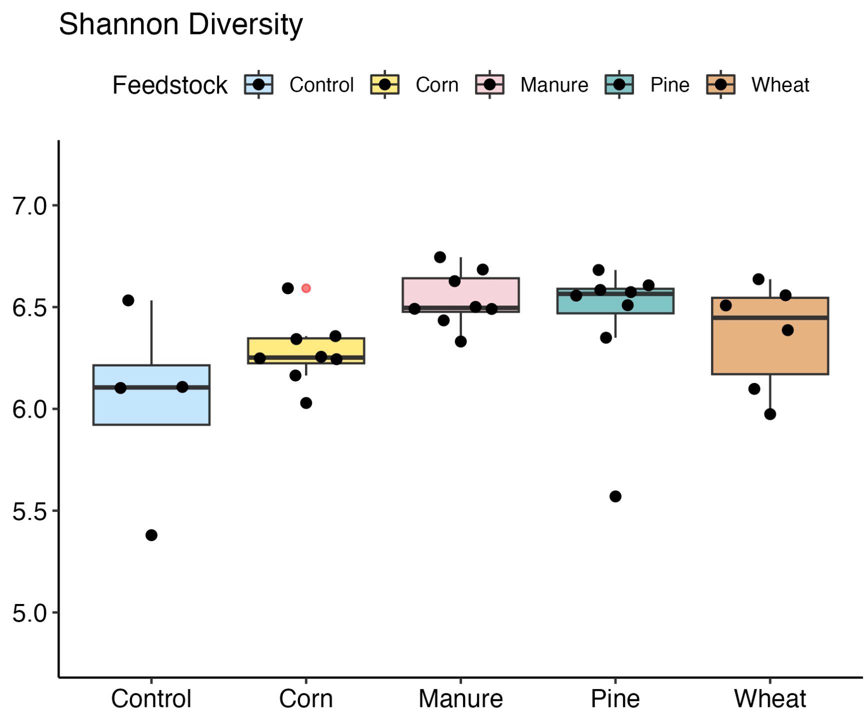


**Fig S3.** Box and whisker plot showing the effects of different types of biochar on the wheat rhizosphere microbiome α diversity. For each biochar type, data from 0.25% and 2.5% application rate were merged. Asterisks indicate significant difference between treatments.

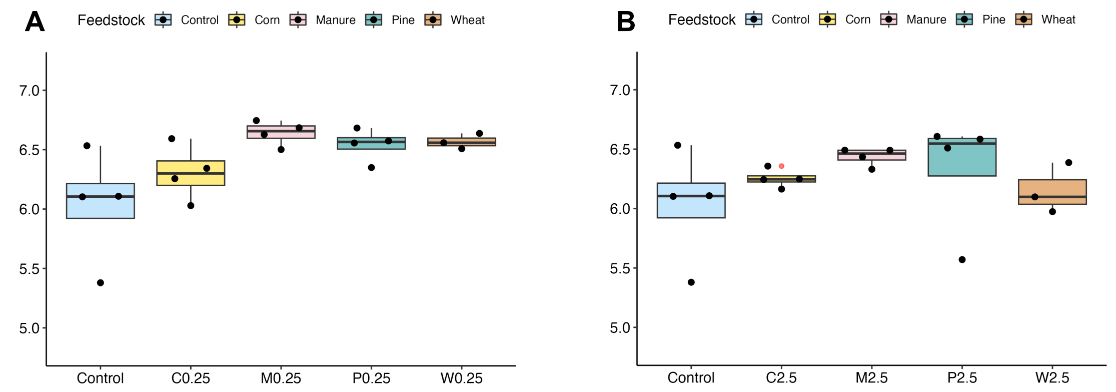


**Fig S4.** Box and whisker plot showing the effects of different types of biochar applied at (A) 0.25% and (B) 2.5% rate on the wheat rhizosphere microbiome α diversity.

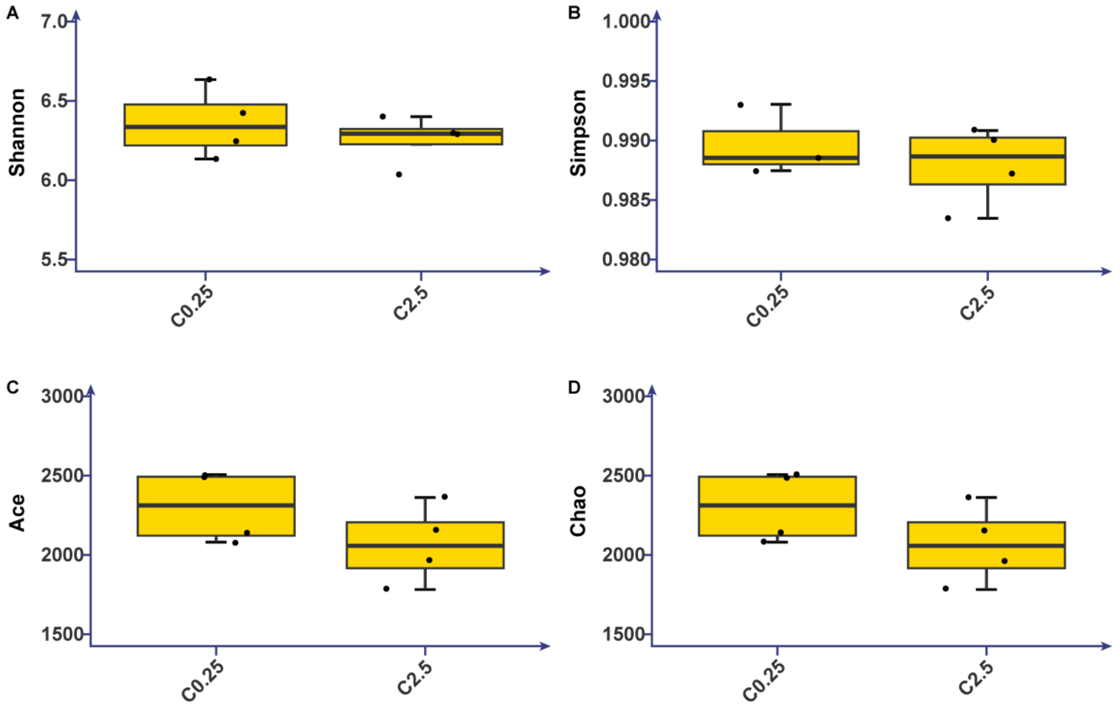


**Fig S5.** Box and whisker plot showing the effects of corn biochar applied at two rates on the wheat rhizosphere microbiome α diversity.

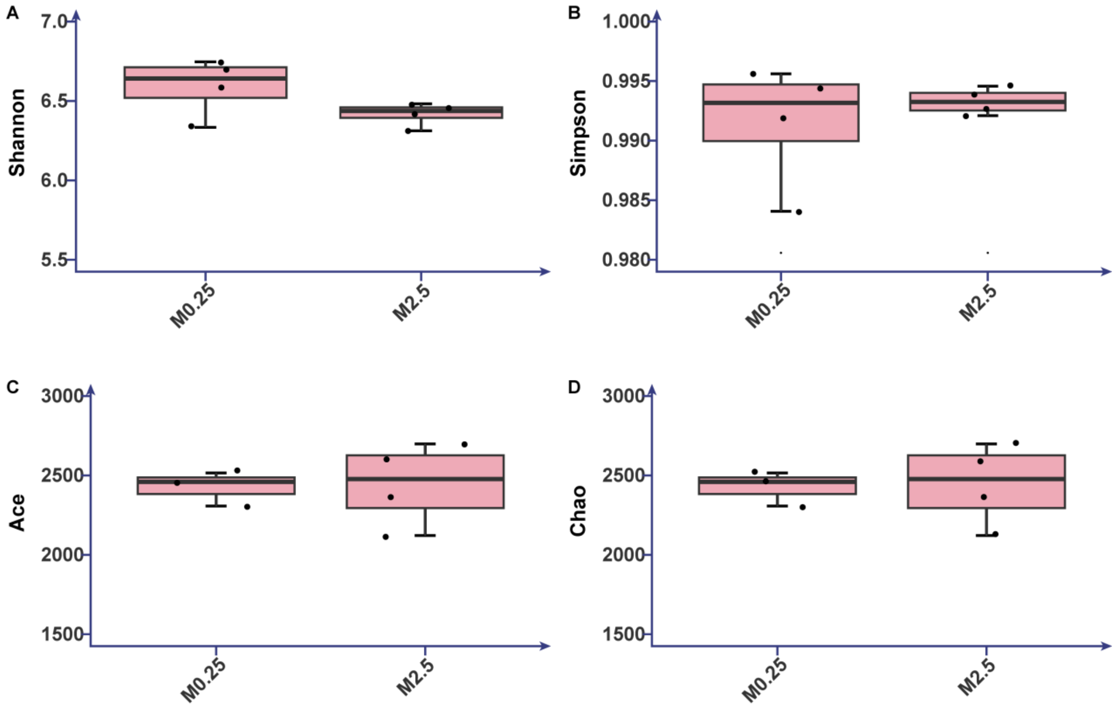


**Fig S6.** Box and whisker plot showing the effects of manure biochar applied at two rates on the wheat rhizosphere microbiome α diversity.

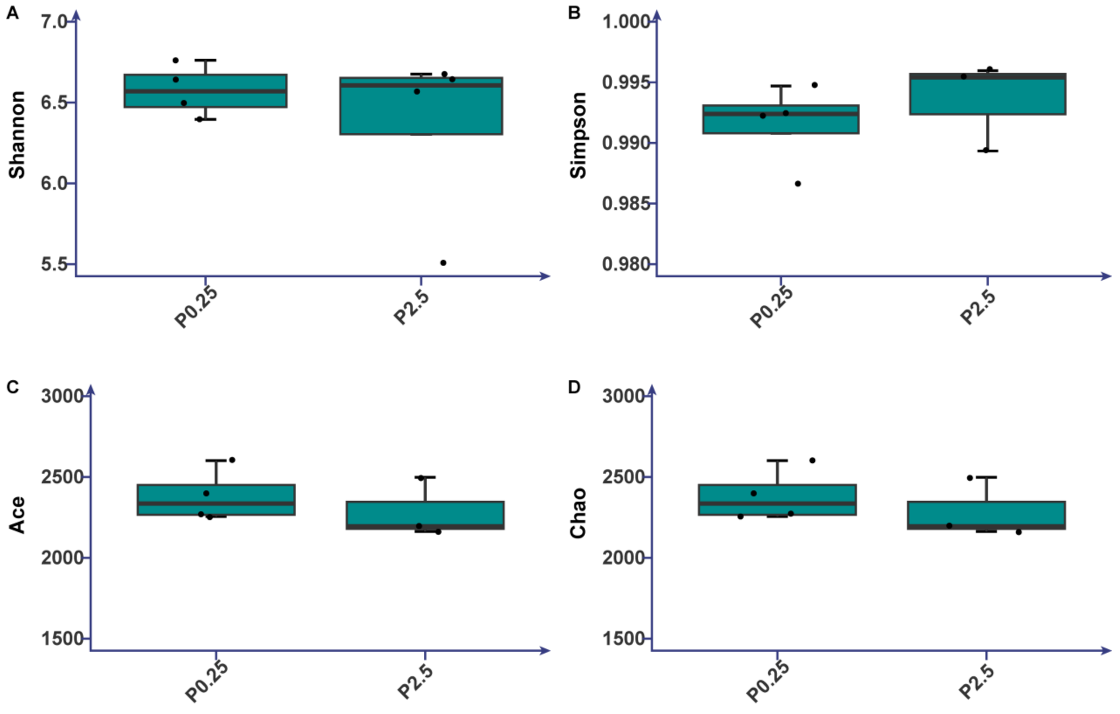


**Fig S7.** Box and whisker plot showing the effects of pine biochar applied at two rates on the wheat rhizosphere microbiome α diversity.

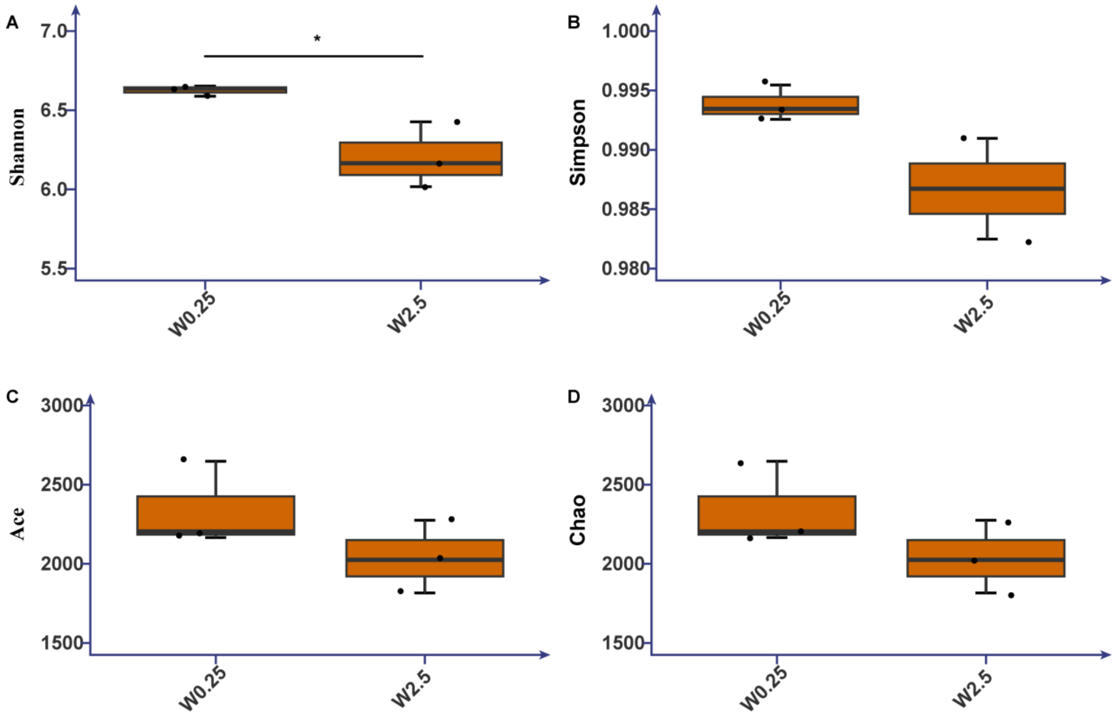


**Fig S8.** Box and whisker plot showing the effects of wheat biochar applied at two rates on the wheat rhizosphere microbiome α diversity. Asterisks indicate significant difference between treatments.

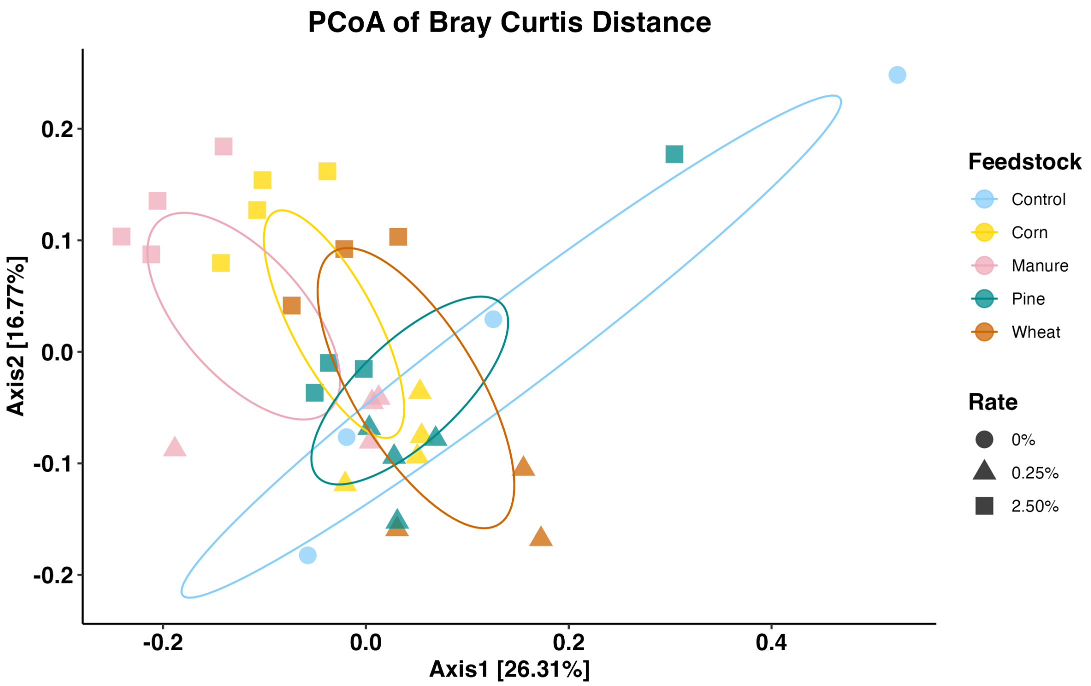


**Fig S9.** PCoA of Bray-Curtis dissimilarity between rhizosphere microbiomes under various biochar treatments.

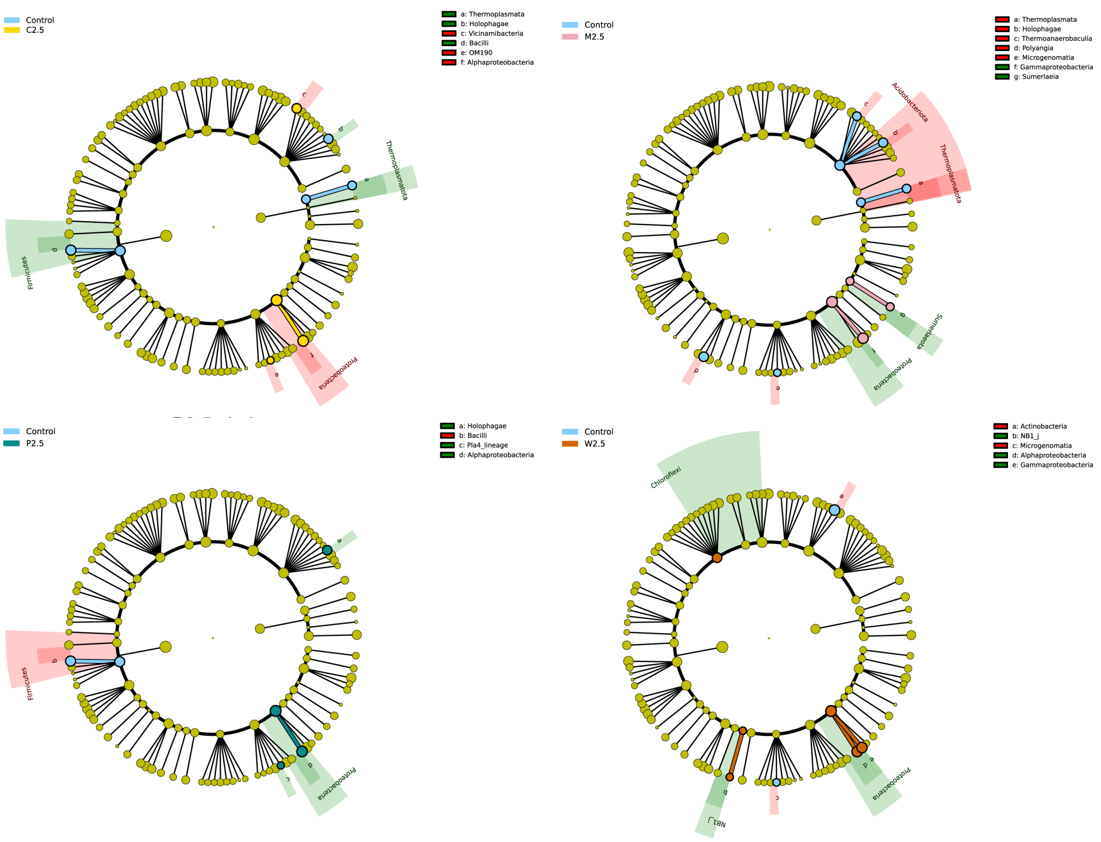


**Fig S10**. Microbial taxa significantly influenced by four types of biochar applied at 2.5% rate. Only phyla and classes are shown in the LEfSe cladograms.

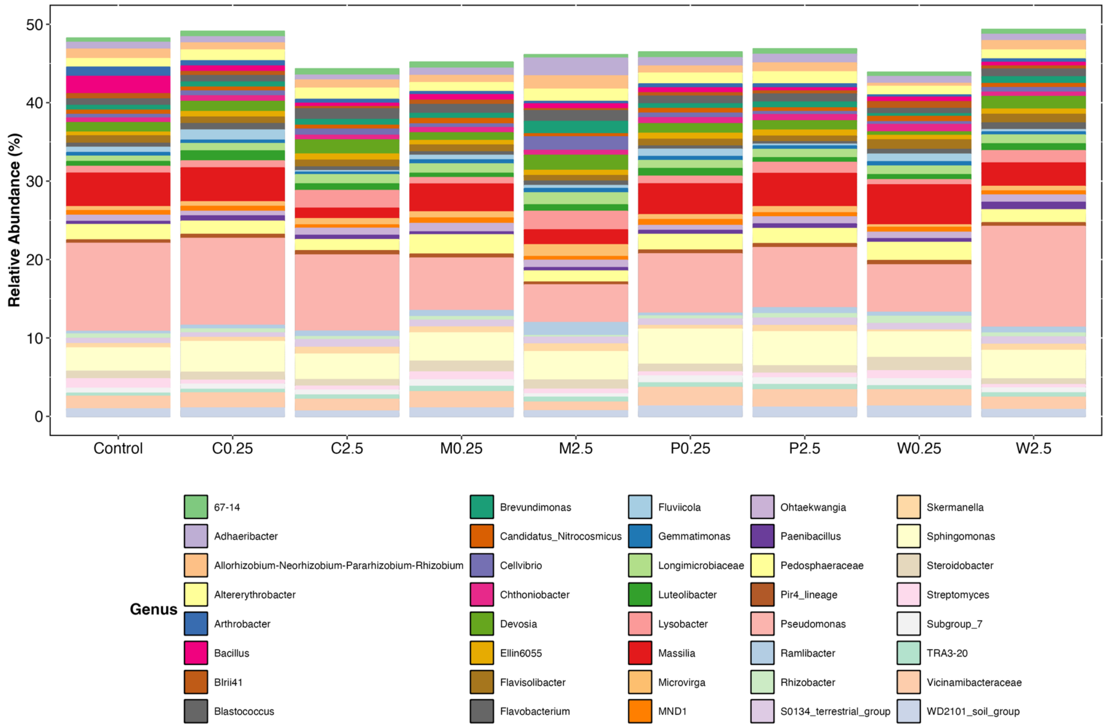


**Fig S11.** Relative abundance of the top 40 rhizosphere microbial genera under various biochar treatments. Data present averages across biological replicates.

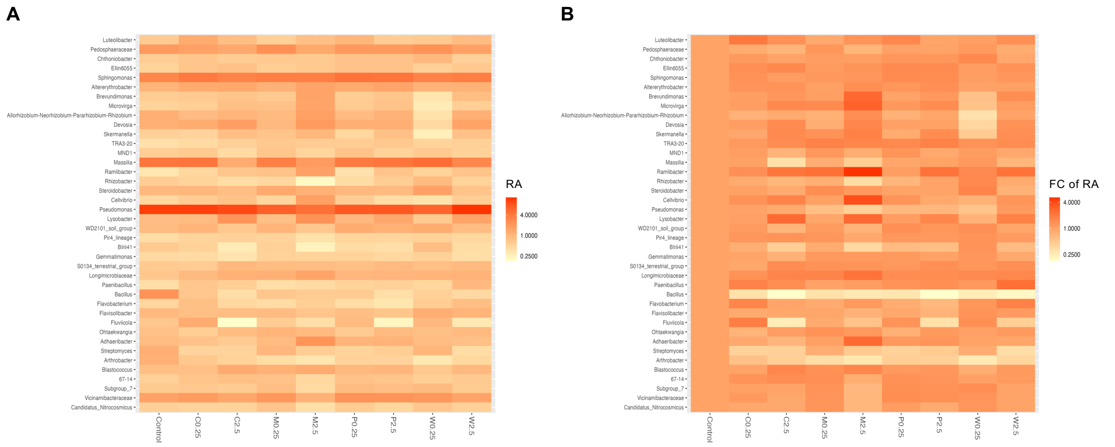


**Fig S12.** Heatmap of (A) relative abundance and (B) fold change of relative abundance of the top 40 rhizosphere microbial genera under various biochar treatments. Data present averages across biological replicates.

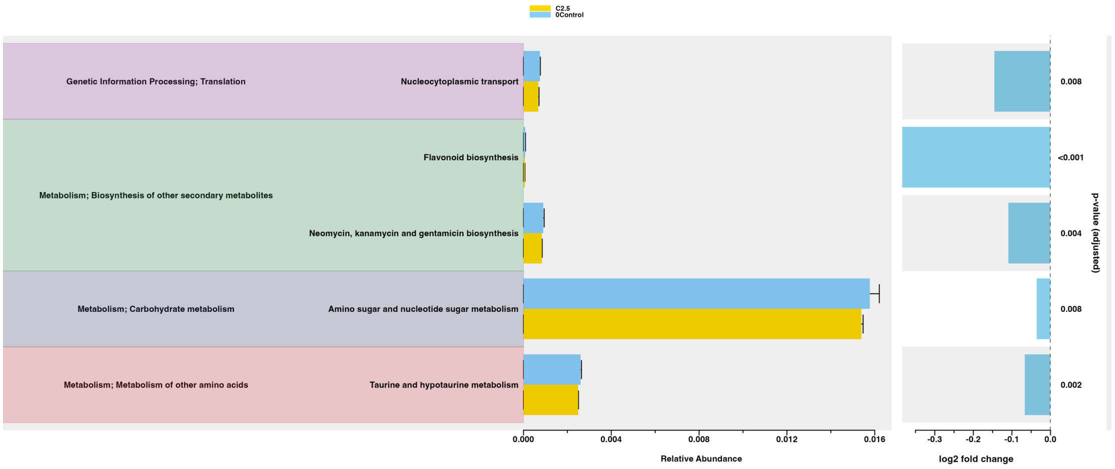


**Fig S13.** Microbial metabolic pathways significantly regulated by C2.5 (DESeq2 test: adjusted-$p < 0.01$). Differences in relative abundance represent averages across biological replicates. Log2 fold change of relative abundance is shown, along with FDR-controlled *p*-values in DESeq2 test. Only one pathway was significantly regulated by C0.25 (Additional file 6).

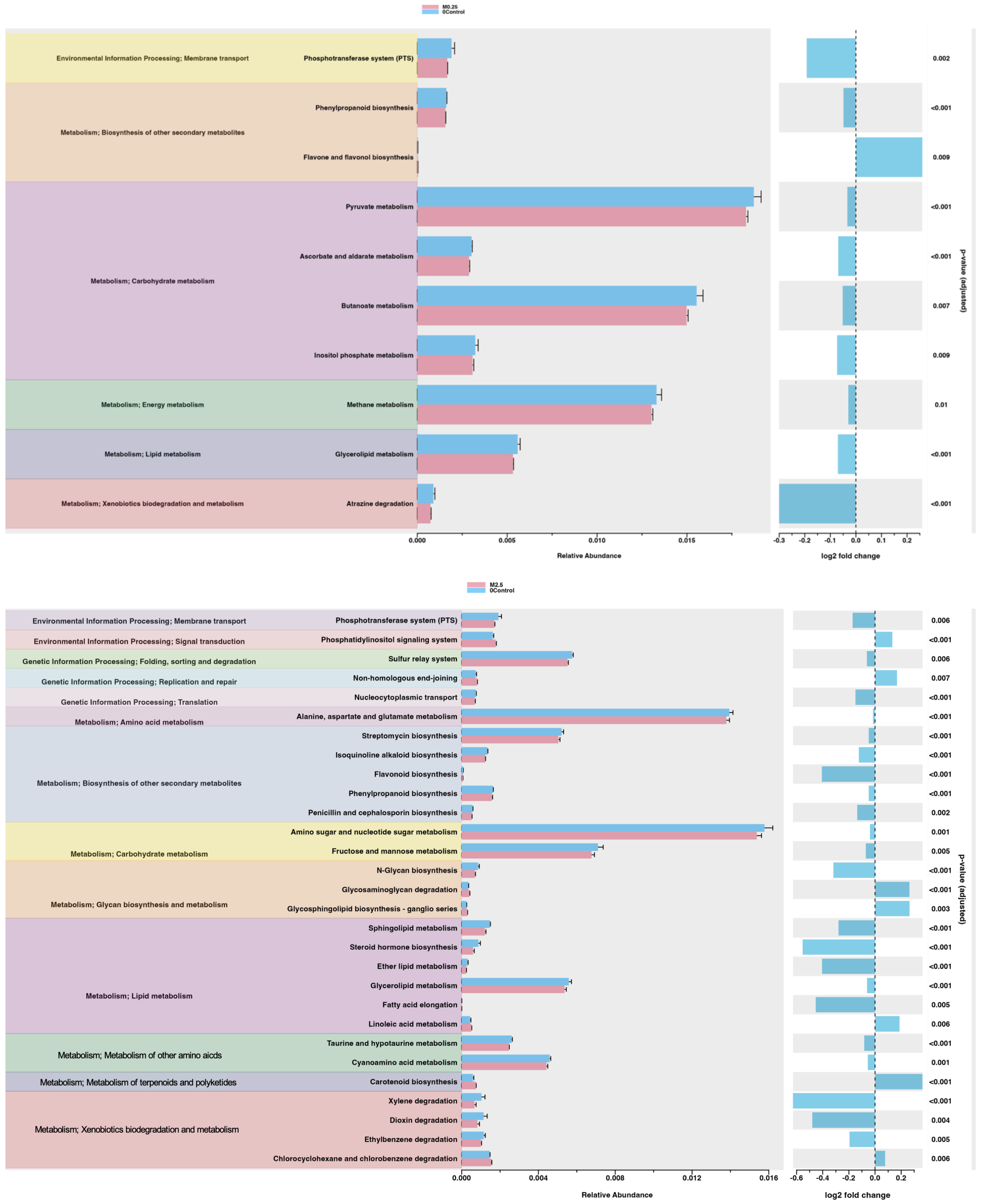


**Fig S14.** Microbial metabolic pathways significantly regulated by (A) M0.25 and (B) M2.5 (DESeq2 test: adjusted-$p < 0.01$). Differences in relative abundance represent averages across biological replicates. Log2 fold change of relative abundance is shown, along with FDR-controlled $p$-values in DESeq2 test.

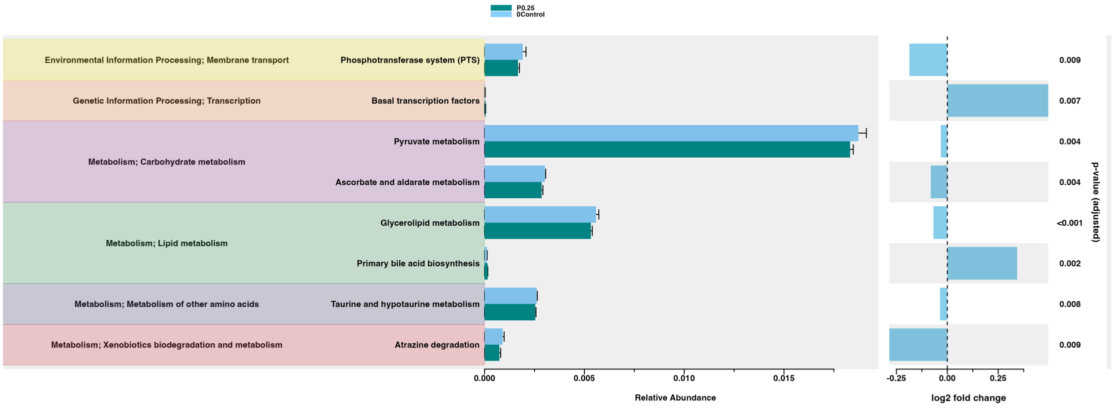


**Fig S15.** Microbial metabolic pathways significantly regulated by P0.25 (DESeq2 test: adjusted-$p < 0.01$). Differences in relative abundance represent averages across biological replicates. Log2 fold change of relative abundance is shown, along with FDR-controlled $p$-values in DESeq2 test. Only one pathway was significantly regulated by P2.5 (Additional file 6).

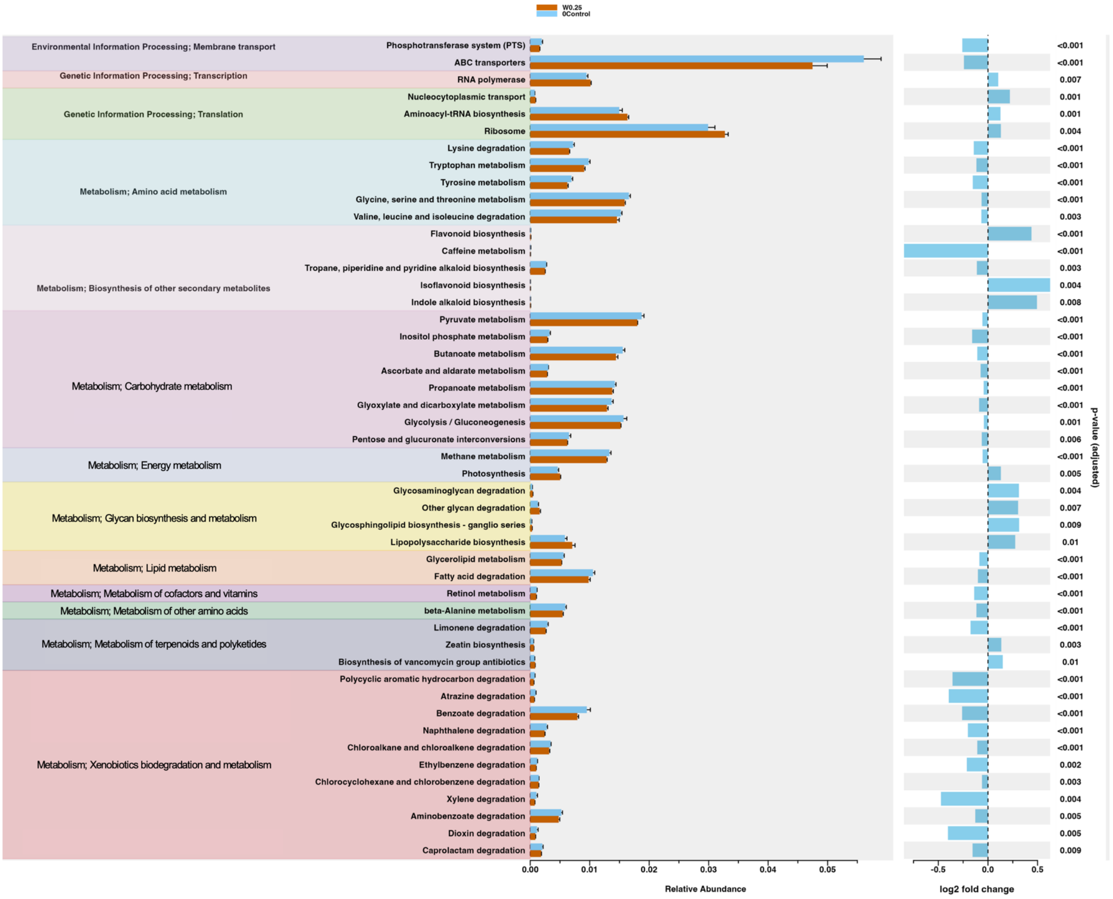


**Fig S16.** Microbial metabolic pathways significantly regulated by W0.25 (DESeq2 test: adjusted-$p$ < 0.01). Differences in relative abundance represent averages across biological replicates. Log2 fold change of relative abundance is shown, along with FDR-controlled $p$-values in DESeq2 test. No pathway was significantly regulated by W2.5 (Additional file 6).

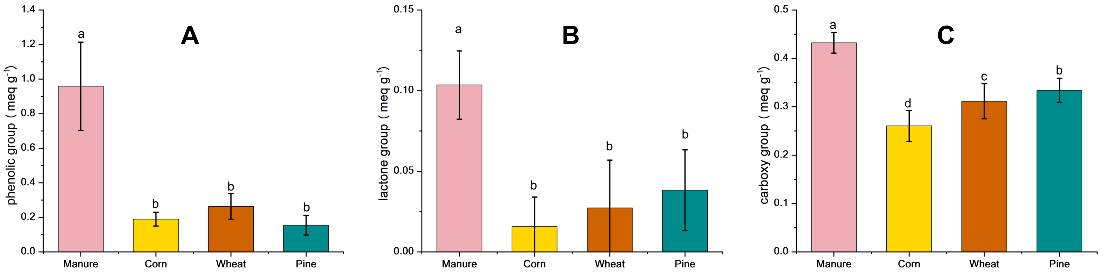


**Fig S17.** Content of (A) acidic carboxyl, (B) lactone, and (C) phenolic group on the surface of different types of biochar. Data represent means and standard deviations from replicate samples.

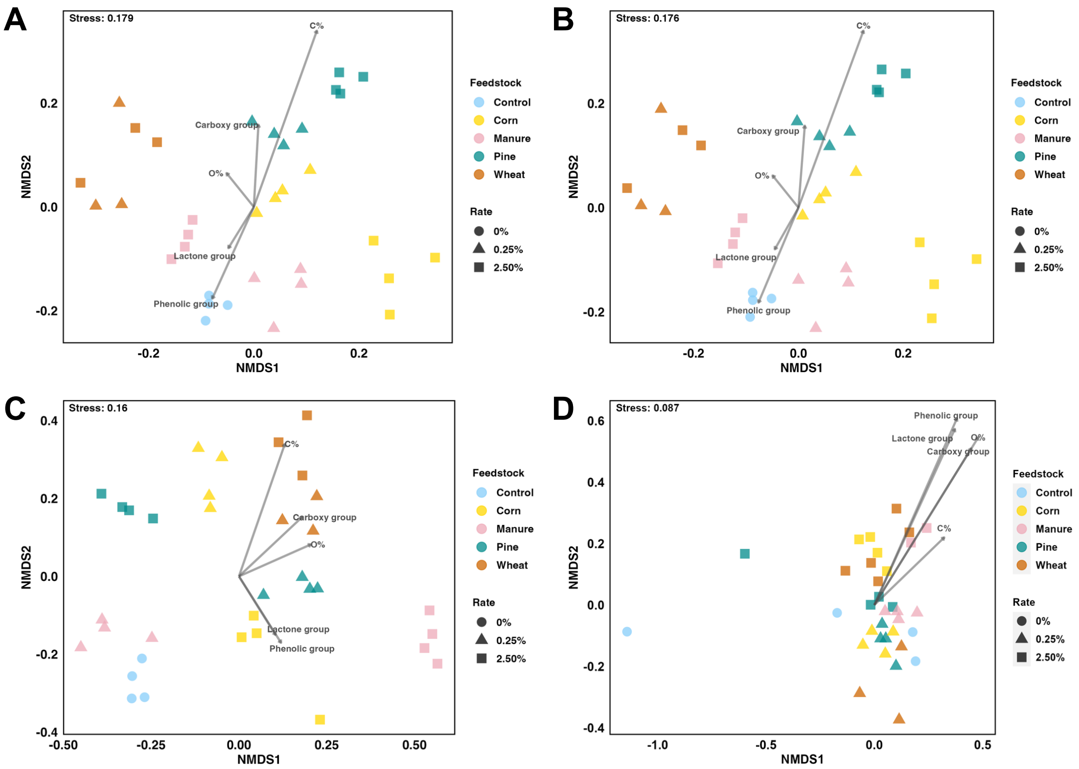


**Fig S18.** Influence of biochar surface C% and O% and three oxygen-containing functional group content on (A) all detected root metabolites, (B) non-glycan metabolites, (C) glycans, and (D) the rhizosphere microbiome. NMDS was based on Bray-Curtis distance. Stress was computed based on 1000 permutations. Feedstock significantly influenced total root metabolites, non-glycan metabolites, and glycans (PERMANOVA: $p < 0.005$). Root glycans were also significantly influenced by biochar application rate (PERMANOVA: $p = 0.01$).

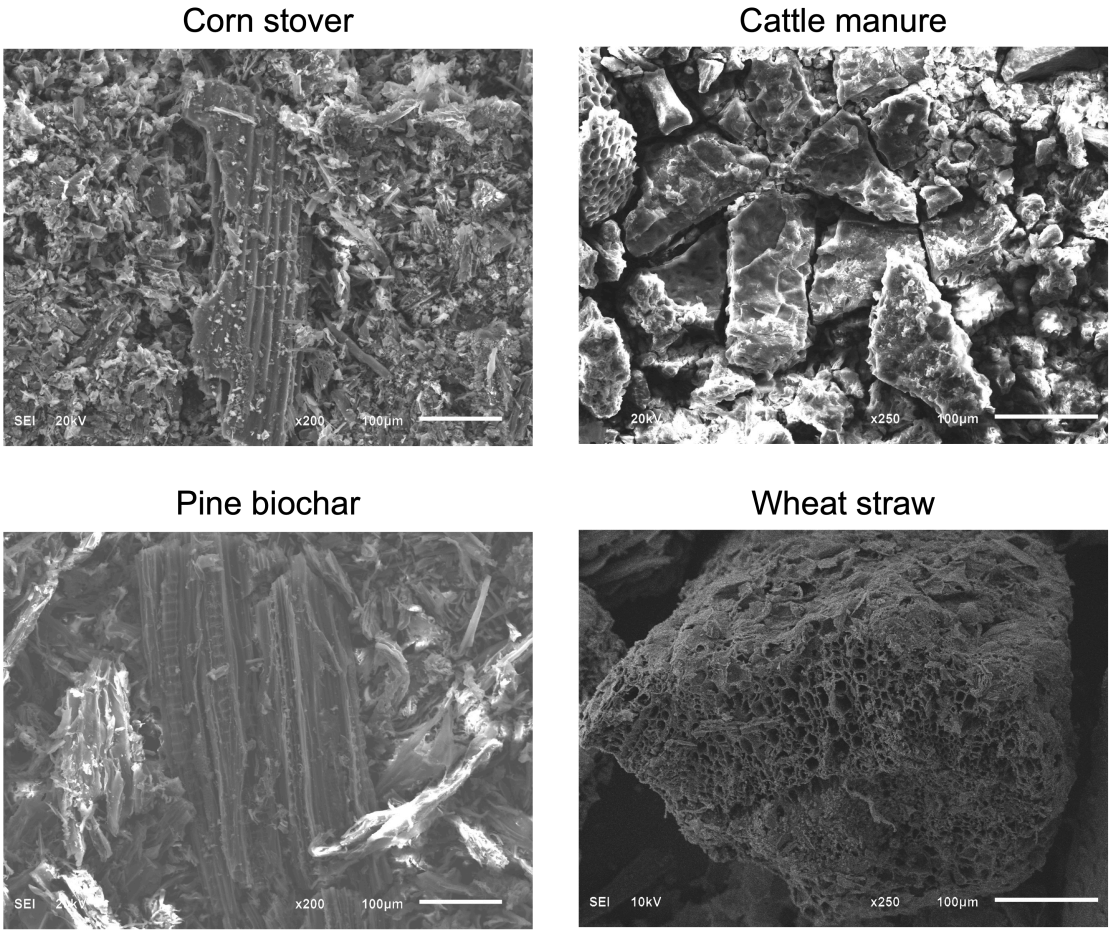


**Fig S19.** Scanning electron microscopic (SEM) images of the four types of biochar used in this study. Biochar was produced using four feedstocks: C, corn stover; M, cattle manure; P, pine sawdust; W, wheat straw.

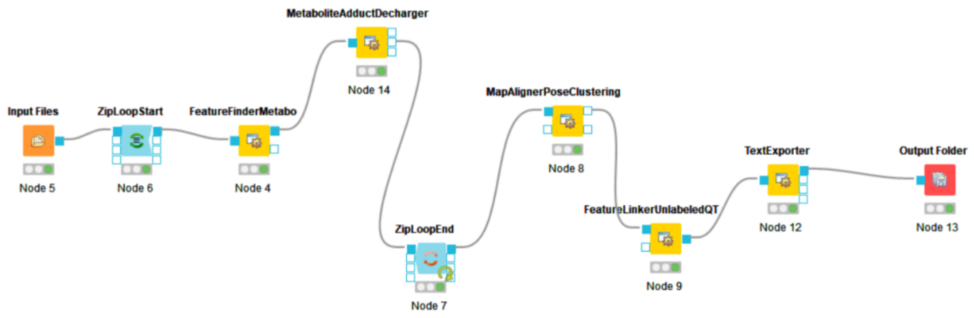


**Fig S20.** Peak identification process in OpenMS platform. This process is different from those in MetaboAnalyst 5.0 and XCMS.
